# Biogenesis of RNA-containing extracellular vesicles at endoplasmic reticulum membrane contact sites

**DOI:** 10.1101/2020.12.04.412379

**Authors:** Bahnisikha Barman, Jie Ping, Evan Krystofiak, Ryan Allen, Nripesh Prasad, Kasey Vickers, James G. Patton, Qi Liu, Alissa M. Weaver

**Author notes:** Current location, Discovery Life Sciences, Huntsville, AL, USA.

## Abstract

RNA transferred via extracellular vesicles (EVs) can influence cell and tissue phenotypes; however, the biogenesis of RNA-containing EVs is poorly understood and even controversial. Here, we identify the conserved endoplasmic reticulum membrane contact site (MCS) linker protein VAP-A as a major regulator of the RNA and RNA-binding protein content of small and large EVs. We also identify a unique subpopulation of secreted small EVs that is highly enriched in RNA and regulated by VAP-A. Functional experiments revealed that VAP-A-regulated EVs are critical for the transfer of miR-100 between cells and for *in vivo* tumor formation. Lipid analysis of VAP-A-knockdown EVs revealed large alterations in lipids known to regulate EV biogenesis, including ceramides and cholesterol. Knockdown of VAP-A-binding ceramide and cholesterol transfer proteins CERT and ORP1L led to similar defects in biogenesis of RNA-containing EVs. We propose that lipid transfer at VAP-A-positive MCS drives biogenesis of a select RNA-containing EV population.

## Introduction

Extracellular vesicles (EVs) are small lipid-bound carriers of bioactive cargoes that are released from diverse cell types to promote cellular communication. Multiple biogenesis mechanisms can promote EV formation and cargo selection, including budding from the plasma membrane as microvesicles and intralumenal budding in endosomes to form exosomes. Recently, it has become apparent that EVs are more heterogeneous than previously appreciated and that diverse cargoes may utilize distinct and potentially non-classical mechanisms for incorporation into EVs (Raposo and Stoorvogel, 2013; van Niel et al., 2018).

In addition to proteins and lipids, EVs contain diverse types of RNA, including miRNAs, lncRNAs, snoRNAs, tRNAs, rRNAs, YRNAs, vault RNAs and mRNAs (Chow et al., 2019; Crescitelli et al., 2013; Driedonks et al., 2018; Hinger et al., 2018; Lasser et al., 2017; Skog et al., 2008; Valadi et al., 2007). These EV-carried RNAs can affect gene expression and the phenotype of recipient cells, which may be important for a variety of diseases (Bell and Taylor, 2017; de Candia et al., 2016; Falcone et al., 2015; O’Brien et al., 2020). EV-enclosed RNA is also being studied for potential use as therapeutics and biomarkers. While extracellular RNA (exRNA) can also be present in a non-vesicular form, encapsulation of RNA in EVs protects it from degradation and allows it to be delivered directly to the cytoplasm of recipient cells via membrane fusion (O’Brien et al., 2020).

Although packaging of exRNAs in EVs is known to be a selective process that leads to the enrichment of certain exRNAs in EVs (Cha et al., 2015; Lee et al., 2019; Lin et al., 2019; Lu et al., 2017), the mechanisms by which this packaging occurs is poorly understood. RNA-binding proteins appear to play a major role in this process, likely controlling both stability and sorting of the RNAs (Deng et al., 2020; Li et al., 2019; McKenzie et al., 2016; Santangelo et al., 2016; Shurtleff et al., 2016; Villarroya-Beltri et al., 2013; Zietzer et al., 2020). We and others identified argonaute 2 (Ago2) and other RISC complex proteins as potent mediators of miRNA sorting into EVs (Bukong et al., 2014; Clancy et al., 2019; Mantel et al., 2016; McKenzie et al., 2016; Melo et al., 2014). Other studies have shown that miRNAs with specific sequence motifs (i.e., GGAG and GGCU) are recognized and selectively sorted into EVs by heterogenous nuclear ribonucleoprotein A2B1 (hnRNPA2B1) and RNA-interacting protein SYNCRIP (Santangelo et al., 2016; Villarroya-Beltri et al., 2013). In addition, the RNA-binding protein Y-box I (YBX-1) is suggested to be involved in the packaging of mRNAs, miRNAs and other exRNAs into EVs by recognizing secondary rather than primary RNA sequence motifs (Kossinova et al., 2017; Shurtleff et al., 2016; Shurtleff et al., 2017; Zietzer et al., 2020).

While certain RBPs are known to be important for determining the RNA content of EVs, it is unclear how these RBP-RNA complexes are trafficked to and selected for incorporation into newly forming EVs at EV biogenesis sites. One clue may come from the typical cellular location of these known RBPs and their activities. hnRNPA2B1 and SYNCRIP are both hnRNPs that mediate RNA processing and translation, with functions in both the nucleus and at the endoplasmic reticulum (ER). Likewise, YB1 affects translation of select mRNAs, localizing to ribosomes and the ER (Matsumoto et al., 2005). Finally, miRNA-loaded Ago2 was shown to be physically associated with rough ER membranes (Barman and Bhattacharyya, 2015; Stalder et al., 2013) before moving to multivesicular bodies for degradation (Bose et al., 2017).

Membrane contact sites with the endoplasmic reticulum (ER MCS) are areas of close apposition between the ER and other organelles (Phillips and Voeltz, 2016; Wu et al., 2018). Key described functions of ER MCS include calcium and lipid exchange between the organelles, and organelle fission. However, the major physiological functions of MCS are still under investigation. A number of tether proteins have been identified that mediate these contacts and control molecular signaling and exchange at these contact points by binding to additional proteins (Phillips and Voeltz, 2016; Wu et al., 2018).

Vesicle-associated membrane protein-associated protein-A (VAP-A) is a major ER MCS tether protein and can undergo both homodimerization and heterodimerization with its homologue VAP-B (Kamemura and Chihara, 2019; Murphy and Levine, 2016). VAP-A interacts with multiple endosome-localized lipid transport proteins, including STARD3, CERT, and ORP1L. ORP1L mediates cholesterol transfer at ER-endosome MCSs (Hanada et al., 2003; Luo et al., 2019; Muallem et al., 2017; Murley et al., 2017; Phillips and Voeltz, 2016; Wu et al., 2018). CERT binds to VAP via its FFAT domain and phosphorylated phosphatidylinositols on other organelles (Kawano et al., 2006). CERT is best known to mediate ceramide transfer from the ER to the Golgi at ER-Golgi MCSs through its StART domain (Hanada et al., 2003; Peretti et al., 2008). Recently, CERT was also shown to mediate ceramide transport at ER-endosome MCS to regulate exosome biogenesis in palmitate-stimulated hepatocytes (Fukushima et al., 2018). Transfer of ceramide via MCS could potentially provide an alternative mechanism to *in situ* ceramide synthesis, which is known to promote nonclassical exosome biogenesis (Trajkovic et al., 2008); however, its general relevance and impact on specific cargoes has not been determined.

To test whether ER MCS are important for biogenesis of RNA-containing EVs, we knocked down (KD) or overexpressed VAP-A in colon cancer cells. We identified multiple small RNAs and RBPs that are differentially altered in both small and large EVs compared to control cells. In-depth analysis of alterations in small EVs revealed that VAP-A promotes formation of a select subpopulation of small EVs that carries the majority of RNA. Moreover, VAP-A-regulated EV biogenesis controls the ability of colon cancer cells to transfer miR-100 to recipient cells and to grow tumors in xenograft mouse experiments. Investigation of the molecular mechanism revealed that VAP-A and its binding partners ORP1L and CERT are important for controlling both the lipid and RNA content of both small and large EVs. These data suggest a model in which lipid transfer at ER MCS drives biogenesis of a select subpopulation of EVs containing RNA-RBP complexes.

## Results

To explore whether the ER may be associated with RBPs trafficked into EVs, we mined publicly available EV proteomics data. Analysis of the human RNA Binding Proteome (Hentze et al., 2018), EV proteome (Kalra et al., 2012; Pathan et al., 2019) and ER proteome (http://www.proteinatlas.orq) (Thul et al., 2017) revealed that 52% (809 RBPs out of 1542 RBPs) of RBPs are secreted in EVs. Among them, 8% are ER–associated proteins (61 RBPs out of 809 RBPs) (Figure 1A). We also examined a recent report in which the most highly represented RBPs across the EV proteomics datasets in the online database EVpedia were manually identified (Mateescu et al., 2017). Of these 80 RBPs, 28% are ER-associated (22 RBPs out of 80 RBPs), and an additional 18% are ribosomal proteins (Figure 1B). Together with previous reports showing that Ago2-miRNA complexes are assembled at ER-associated ribosomes (Maroney et al., 2006; Nottrott et al., 2006), these data led us to hypothesize that a significant portion of EV-incorporated RNAs and RBPs are associated with the ER.

**Figure 1:**
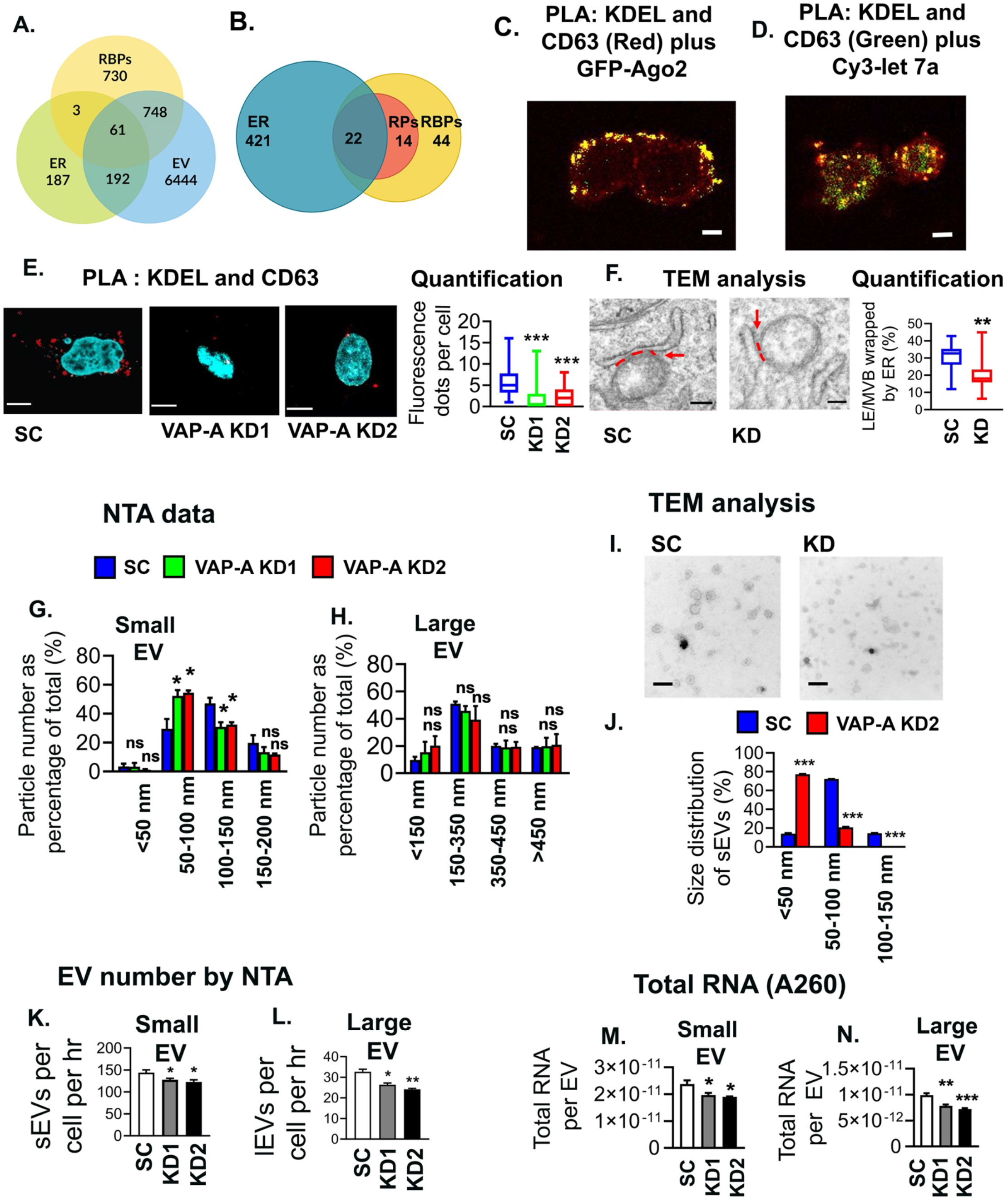
ER membrane contact sites and VAP-A are associated with biogenesis of RNA-containing EVs. (A) Venn diagram shows the overlap of human RBPs (1542) (Hentze et al., 2018), endoplasmic reticulum (ER) proteins (443) (Thul et al., 2017), and extracellular vesicle (EV) proteins (7445) (from Vesiclepedia (Kalra et al., 2012; Pathan et al., 2019)). (B) Representation of previously published top 80 EV associated RBPs (Mateescu et al., 2017) present on ER membrane. Venn diagram shows 22 RBPs (28%) are ER associated and an additional 14 RBPS (18%) are ribosomal proteins (RPs). (C) Representative merged image for proximity ligation assay (PLA) analysis for ER MCS (red) in GFP-Ago2 (green) expressing DKs-8 cells. PLA reaction was performed with KDEL (ER marker) and CD63 (late endosome/MVB marker) and appears as red fluorescent dots. DAPI (blue) was used to stain the nucleus. Yellow indicates colocalization of Ago2 with ER MCS. (D) Representative merged image for PLA analysis of ER MCS (KDEL+CD63 = green) in cy3-let-7a (red) expressing DKs-8 cells. DAPI (blue) was used to stain the nucleus. (E) Representative merged images of PLA analysis of ER MCS (KDEL+CD63, red) in control (Sc) and VAP-A KD1 and KD2 DKs-8 cells. DAPI, blue. Fluorescence dots per cell were calculated and plotted from sixty cells in three independent experiments. Box and whiskers plots with box indicating 25th-75^th^ percentile, whiskers showing 5-95^th^ percentile, and line indicating median. (F) Representative TEM images of control (Sc) and VAP-A KD2 (KD) DKs-8 cells. ER-endosome MCS are indicated by a red dashed line and arrow. Quantification shows area of LE/MVB membrane in contact with ER and plotted as percentage of the endosome circumference. Box and whiskers plots with box indicating 25th-75^th^ percentile, whiskers showing 5-95^th^ percentile, and line indicating median. (G and H) Graphs show quantitation from nanoparticle tracking analyses (NTA) of small and large EVs isolated from control (Sc) and KD DKs-8 cells. Particle numbers were plotted according to their size from three independent experiments. Note a shift in the KD small EV population towards smaller sizes. (I and J) Representative TEM images and size analysis for small EVs purified from DKs-8 control (Sc) and VAP-A KD2 (KD) cells. Quantification of 150 vesicles total per condition (control or KD) from three independent experiments. (K and L) Graphs of EV release rate from control and KD cells quantitated from NTA data and normalized based on final cell number and conditioned media collection time. Data from five independent experiments. (M and N) Graphs of total RNA concentration measured by NanoDrop (A260) for small and large EVs isolated from control (Sc) and VAP-A KD DKs-8 cells. Data from five independent experiments. Bar graphs indicate mean +/- S.E.M. *p<0.05, **p<0.01, ***p<0.001. ns, not significant. See also Figures S1 and S2.

To assess whether RNA and RBPs known to be selectively incorporated into EVs localize to ER MCS, we identified ER-endosome MCS with a proximity ligation assay using antibodies against KDEL and CD63 (Figure 1C) in DKs-8 colorectal carcinoma cells. These cells were previously reported to export RBPs and RNAs in EVs, including Ago2 and let7a miRNA (Demory Beckler et al., 2013; McKenzie et al., 2016). We localized either GFP-Ago2 or Cy3-let7a to the identified MCS and found that both had significant overlap with MCS (Figures 1C, 1D, S1A).

### The MCS tether protein VAP-A controls the size and cargo content of EVs

To determine whether ER MCS may affect the biogenesis of RNA-containing EVs, we knocked down (KD) the ER MCS linker protein VAP-A in DKs-8 colon cancer cells. (Figures S1B). Both proximity ligation and transmission electron microscopy (TEM) analyses confirmed a reduction in ER-endosome MCS in VAP-A KD cells compared with controls (Figures 1E, 1F). By contrast, loss of VAP-A had no discernable effect on cell viability or ER stress markers, suggesting that loss of VAP-A did not generally disrupt cell functions (Figures S1C-S1F). Nanoparticle tracking analysis of EVs purified by cushion density gradient (Li et al., 2018) (Fig S1G, S1H) from DKs-8 control or VAP-A KD cells revealed a decrease in the number and size of small EVs and in the number of large EVs purified from VAP-A KD cells (Figures 1G, 1H, 1K, 1L). The alterations in small EV size with VAP-A KD were further validated by analysis of TEM images of negatively stained small EVs (Figures 1I and 1J). We found similar alterations in size and number of EVs purified from control and VAP-A KD DKs-8 cells for a second colon cancer cell line, DKO-1 (Figures S2A-S2E). Western blot analysis of our small and large EV preparations confirmed the presence of typical EV marker proteins and the absence of the negative marker GM130 (Thery et al., 2018) (Figure S1G).

To determine whether VAP-A affects the RNA content of EVs, total RNA was extracted and analyzed. Assessment of the total RNA content of small and large EVs by A260 reading with a NanoDrop indicated that VAP-A KD EVs contained significantly less RNA than control EVs (Figures 1M and N). To more comprehensively assess the effect of VAP-A on EV RNA content and assess whether certain RNAs are more dependent on MCS for trafficking into EVs, we performed next generation sequencing on equal amounts of small RNA purified from control and VAP-A KD cells and EVs (Supplementary Datasheets 1-3). Principal component analysis of the data revealed that VAP-A KD alters the small RNA profiles of small EVs, large EVs, and cells (graphs for miRNA shown in Figures 2A, S3A and S3B). To identify individual miRNAs whose secretion was altered by VAP-A, we normalized the miRNA levels in EVs to the levels in the corresponding cells of origin. Using a criterion of ≤ 0.5 or ≥ 2 fold change and FDR ≤ 0.05, we identified 82 miRNAs that were differentially exported into VAP-A KD EVs compared to control EVs (Figure 2B). Of these, 26 were common to both small EVs and large EVs (Figure 2C). To validate our sequencing results, we performed qRT-PCR analysis for specific miRNAs taken from our sequencing dataset (miR-371a, miR-372) that were downregulated in VAP-A KD EVs. We also analyzed 4 miRNAs known to be selectively exported in DKs-8 EVs: let-7a, miR-100, miR-320, miR-125b ((McKenzie et al., 2016) and unpublished data). These miRNAs were also decreased in KD EVs in our dataset but did not reach the criteria of FDR ≤ 0.05. The levels of candidate RNAs were normalized to U6, which is exported in EVs but not affected by VAP-A. We found that all of the candidate miRNAs were decreased in both small and large EVs (Figures 2D-2E). Consistent with a specific role for VAP-A in RNA sorting into EVs, there was either no change or an increase in the cell levels of the same RNAs (Fig 2F).

**Figure 2:**
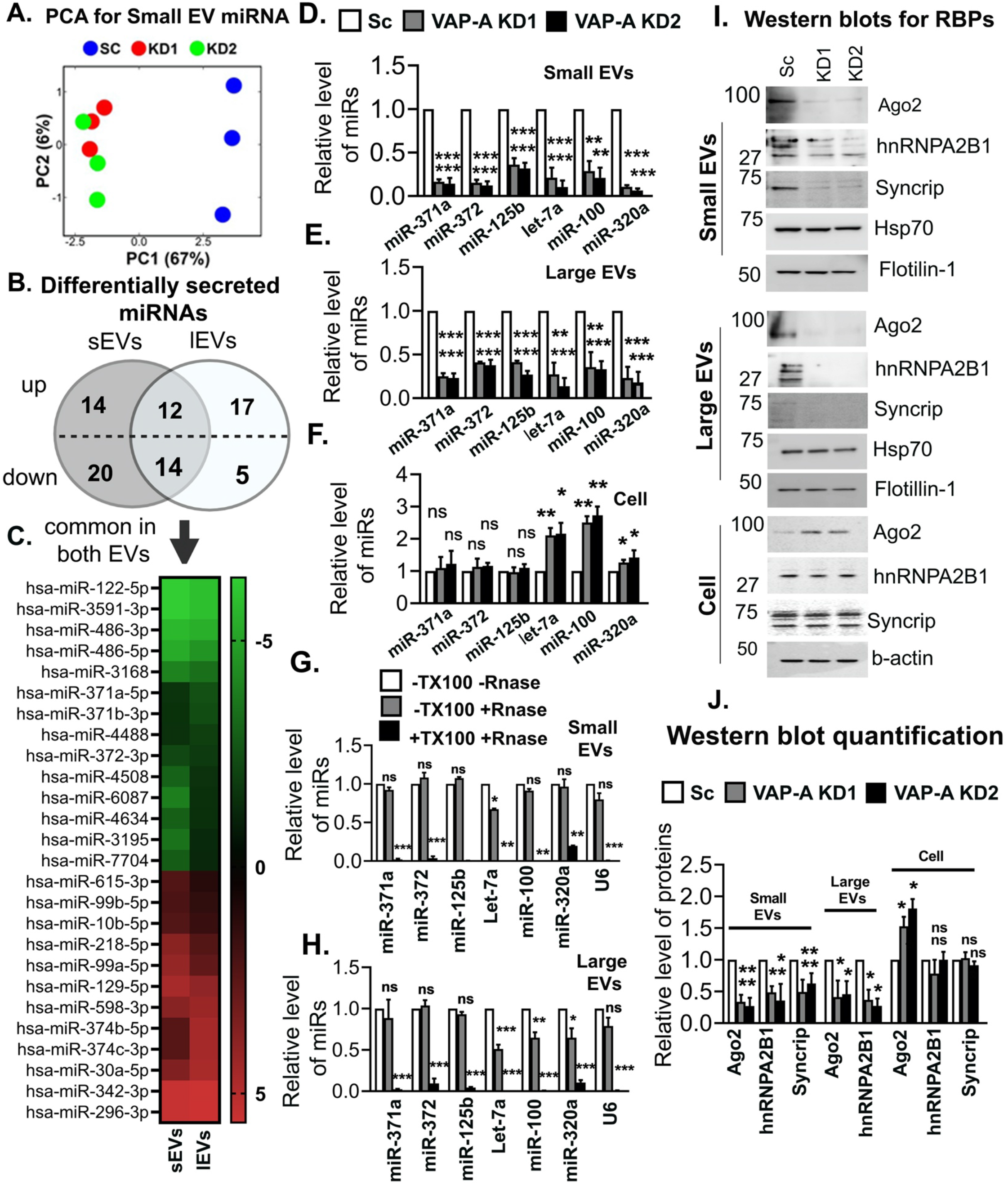
VAP-A regulates the miRNA composition of small and large EVs. (A) Principal component analysis (PCA) of miRNA composition in small EVs showing segregation of KD (KD1 and KD2) from control (SC) data. (B) Venn diagram depicts numbers of up-or down-regulated miRNAs in small and large EVs upon VAP-A knockdown. 2-fold changes of miRNAs in small and large EVs were normalized by levels in their parental cells. miRNAs were considered significantly changed if either ≥ 2-fold or ≤ 0.5-fold enriched with a FDR value ≤ 0.05. (C) Heat map represents levels (log 2-fold change) of differentially secreted 29 miRNAs in small and large EVs compared to their parental cells upon VAP-A KD. (D-F) Relative levels of miR-371a, miR-372, miR-125b, let-7a, miR-100, miR-320a in control and VAP-A KD small EVs, large EVs and cells. Quantitative RT PCR was performed with 10 ng of total RNA. All experiments were done in triplicate. U6 snRNA was used to normalize Ct values. (G-H) Relative levels of miRNAs (miR-371a, miR-372, miR-125b, let-7a, miR-100, miR-320a) in small and large EVs purified from control DKs-8 cells. Small and large EVs were treated with (+) or without (-) RNase in absence (-) or presence (+) of 1% Triton-X-100 (TX100) followed by total RNA isolation and qRT-PCR. All experiments were done in triplicate. (I and J) Immunoblots show levels of RBPs (Ago2, hnRNPA2B1, Syncrip) and EV markers (flotillin-1, HSP70) in control (Sc) and VAP-KD EVs and cells. Quantification of immunoblot from three independent experiments were shown. Values were normalized by HSP70 (for small and large EVs) and by beta-actin (for cell lysates). Data were plotted as mean +/- S.E.M. *p<0.05, **p<0.01, ***p<0.001. ns, not significant. See also Figures S2 and S3.

We also analyzed snoRNA levels in EVs and cells from our RNA-seq dataset. Using our previous criteria, we found alterations in secretion of 14 snoRNAs (11 reduced and 3 increased) in small EVs with VAP-A KD, but no alterations in snoRNAs in large EVs (Fig S3C). qRT-PCR for specific snoRNAs (snoRD105, snoRA40, snoRA42 and snoRD45) taken from our dataset revealed that all of the snoRNAs were reduced in VAP-A KD small EVs but unchanged in KD cells (Figs S3D and S3F). In addition, snoRA42 and snoRD45 levels were reduced slightly in VAP-A KD large EVs (Fig S3E). Analysis of the RNA-Seq dataset also revealed alterations in the levels of tRNA fragments in VAP-A KD EVs (Figures S3G and S3H).

To further test our hypothesis that VAP-A is a positive regulator of EV number and cargo content, we overexpressed VAP-A (Fig S4A). Consistent with that hypothesis, we found that overexpression of VAP-A in DKs-8 cells increased the number of small and large EVs per cell, the total level of RNA per EV, and the levels of specific miRNAs in small and large EVs (Figures S4B-S4K). Interestingly, the levels of those same miRNAs in VAP-A-OE cells were significantly decreased, suggesting that export of miRNAs into EVs may impact their levels in cells (Chiou et al., 2018).

Since non-vesicular RNAs can associate with the outside of EVs in a nonspecific manner and could theoretically contaminate our assays, we analyzed whether the miRNAs associated with our EVs were sensitive to RNase in the absence or presence of detergent. We found that the six candidate miRNAs and U6 are only minimally depleted by RNase treatment in the absence of detergent but are almost fully depleted by RNase in the presence of detergent (Figures 2G and 2H). These data are consistent with the candidate RNAs being on the inside of the EVs, as would be expected by a selective biogenesis mechanism.

Previous reports have shown that RBPs such as Ago2, hnRNPA2B1, and SYNCRIP, are involved in RNA sorting to EVs (McKenzie et al., 2016; Santangelo et al., 2016; Villarroya-Beltri et al., 2013). Western blot analysis revealed that Ago2 and hnRNPA2B1 are reduced in both small and large EVs isolated from VAP-A KD cells while SYNCRIP is reduced in small EVs from KD cells and undetectable in large EVs (Figures 2I and 2J). To test whether the RBPs we detect in our Western blots are on the inside or outside of EVs in our preparations, we used a previously published dot blot method (Lai et al., 2015; McKenzie et al., 2016; Sung and Weaver, 2017). Serially diluted EV samples were dotted onto nitrocellulose membranes and immunoblotted for Ago2, hnRNPA2B1, CD63, or flotillin in the presence or absence of 0.1% Tween-20 detergent to permeabilize the EVs. As the antibody to CD63 was to an extracellular epitope, it served as a positive control for small EVs and was detected in both the presence and absence of detergent (Fig S5A). Flotillin-1 was used as a control for large EVs, as they do not have detectable CD63 (Fig S1C). As expected, flotillin-1 was mostly detected on the inside of EVs (Fig S5B, not that it is possible that some membrane flipping may occur for microvesicles (Beer et al., 2018)). By contrast, Ago2 and hnRNPA2B1 were detected only in the presence of detergent, indicating that they are present inside the vesicles (Figures S5A and S5B) and unlikely to represent protein aggregate contamination of our EV preparations.

### A subpopulation of small EVs contains the majority of RNA and is regulated by VAP-A

A central question in the field has been whether RNA is contained in a subset of EVs or is uniformly distributed at low levels in most EVs (Chevillet et al., 2014). To address this question and determine whether VAP-A regulates biogenesis of a subset of cellular EVs containing RNAs, we used a previously published density gradient protocol (Kowal et al., 2016) to isolate “light” and “dense” subpopulations of small EVs from control and VAP-A KD cells. Consistent with the previous publication, we found two peaks of EVs on the density gradient, a peak at fraction 3 that represents less dense material and is enriched for the EV markers TSG101 and CD63 and a peak at fraction 5 that contains more dense material and is enriched for EV cargoes Ago2 and fibronectin (McKenzie et al., 2016; Sung et al., 2015) (Figure 3A). Nanoparticle tracking analysis revealed that the majority of the EVs are found in the light fraction (Figures 3B and 3C, compare the Y-axis scales). In addition, VAP-A KD led to a reduction in the number of dense EVs secreted over time but no change in the number of light EVs. In addition, the KD EV population contained more small EVs and fewer large EVs when compared to the control EV population, especially for the dense EV populations (Figures 3D and E). Estimation of the total RNA found in each EV population revealed that the dense EVs are relatively enriched in RNA, compared to light EVs (Figures 3F-I, 6.3-fold and 11.3-fold enrichment comparing control dense to light EVs by NanoDrop and Qubit methods, respectively). qRT-PCR analysis further revealed that VAP-A KD selectively decreases the levels of six miRNAs in dense but not light small EVs (Figure 3J). As with our other method of EV preparation (Fig 2), candidate small RNAs appear to be contained on the inside of both light and dense small EVs, as they are depleted by RNase treatment only in the presence of detergent (Figures 3K and L).

**Fig 3:**
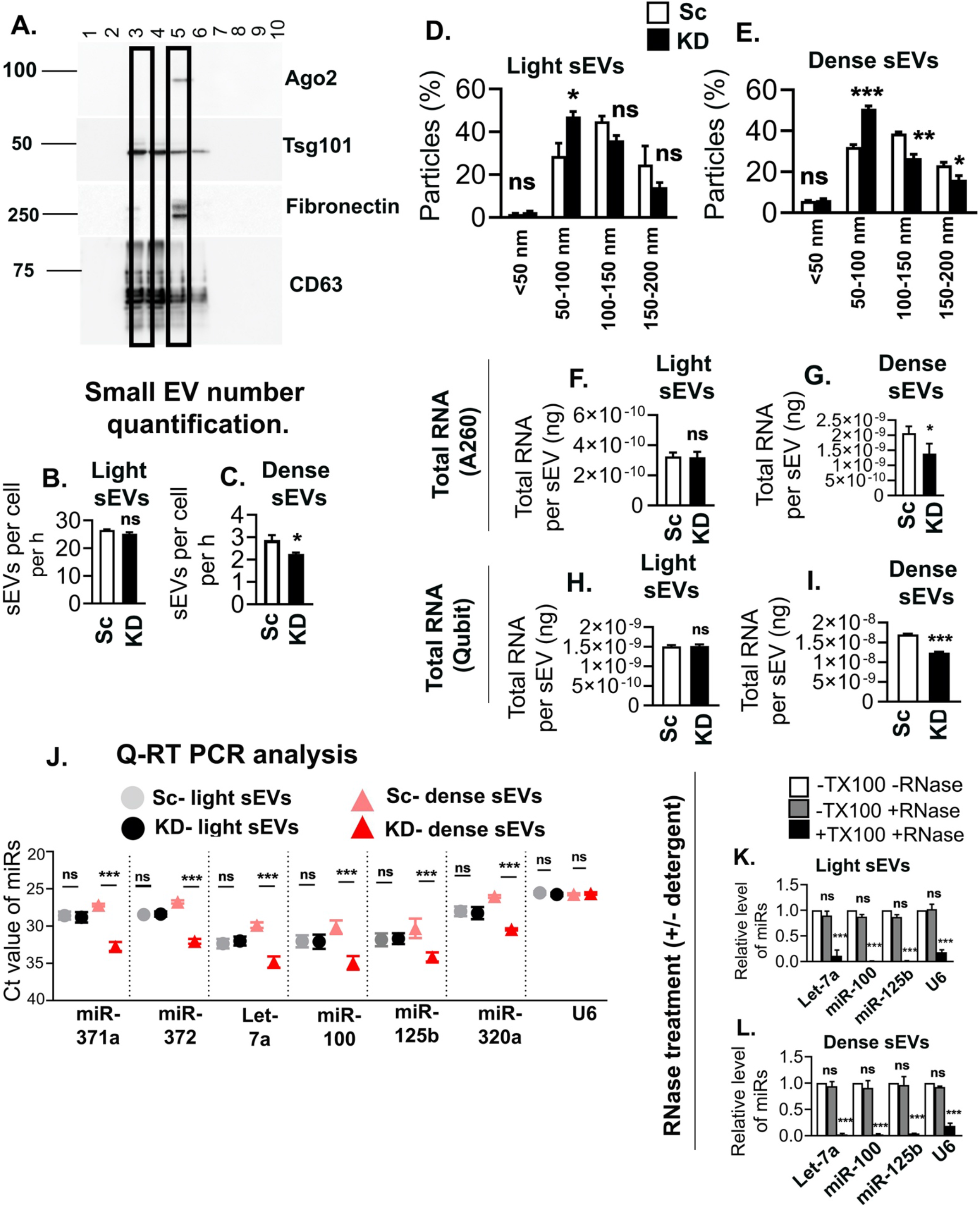
A subpopulation of small EVs is highly enriched in RNA and is regulated by VAP-A. (A) Representative immunoblot of different EV cargo and marker proteins (fibronectin, ago2, Tsg101 and CD63) shows segregation in fractions 3 and 5 of “light” and “dense” EVs purified from control DKs-8 cells. (B and C) Graphs show concentrations of “light” and “dense” EVs purified from control (Sc) and VAP-A KD2 (KD) DKs-8 cells. Data from three independent experiments. Note the difference in the scale of the Y-axes on the two graphs. (D and E). Graphs of NTA data show size distribution of light and dense small EVs isolated from control (Sc) and VAP-A KD2 (KD) DKs-8 cells. Data from three independent experiments. (F - I) Total RNA quantity measured by NanoDrop (A260) (F, G) or Qubit (fluorescence) (H, I) per “light” and “dense” small EV isolated from control and KD cells. Data from three independent experiments. Note the enrichment of RNA in dense EVs compared to light EVs, regardless of the method of measurement. (J) Ct values of specific miRNAs in control and KD light and dense small EVs show VAP-A KD selectively affects the dense EV population. Data from three independent experiments. (K and L) Specific miRNAs (Let-7a, miR-100, miR-320a and U6) are present inside light and dense small EVs, as they are only susceptible to RNase treatment in the presence of Triton-X-100 (TX100). Data from three independent experiments. Data plotted as mean +/- S.E.M. *p<0.05, **p<0.01, ***p<0.001. ns, not significant.

### VAP-A expression controls the function of EVs

To test whether VAP-A affects the function of EVs, we leveraged our previous work, in which we showed that miR-100 can be transferred in a coculture from donor cells grown on Transwell filters to recipient cells present in culture wells below (Cha et al., 2015). Since mutant KRAS-expressing DKO-1 cells secrete more miR-100 in SEVs compared to matched isogenic wild type KRAS DKs-8 cells (Cha et al., 2015), we used control and VAP-A KD DKO-1 cells (Fig S2) as donor cells.

To perform the Transwell co-culture assay, DKs-8 recipient cells were seeded in culture wells and transiently transfected with luciferase reporters containing either 3 artificial miR-100 binding sites in the 3’ UTR (luc-miR-100-PT) or control scrambled sites (luc-con) (Cha et al., 2015) (Figure 4A). Scrambled control (Sc) or VAP-A KD DKO1 cells, or parental DKs-8 cells were used as donors. Consistent with our previous data (Cha et al., 2015), luciferase expression from the miR-100-PT reporter was significantly reduced in recipient DKs-8 cells when co-cultured with control DKO-1 cells in comparison to either the DKs-8 donor cells or the no donor condition (Figure 4B). VAP-A KD in DKO-1 donor cells reversed this reduction in luciferase, bringing it back to the levels found in the no donor or Dks-8 donor conditions. There were no alterations in luciferase expression from the control reporter under any of the conditions (Figure 4B). To confirm that the effects of VAP-A KD in the coculture system were due to EV transfer, we purified small EVs from control or VAP-A KD DKO-1 cells or from DKs-8 cells. As expected, control DKO-1 EVs contained ~2-fold more miR-100 than did KD DKO-1 EVs or DKs-8 EVs (Figure 4C). When added to donor cells expressing miR-100-PT luciferase, the control DKO-1 EVs, but not the VAP-A KD EVs, reduced luciferase expression similar to the co-culture results (Figure 4D).

**Figure 4:**
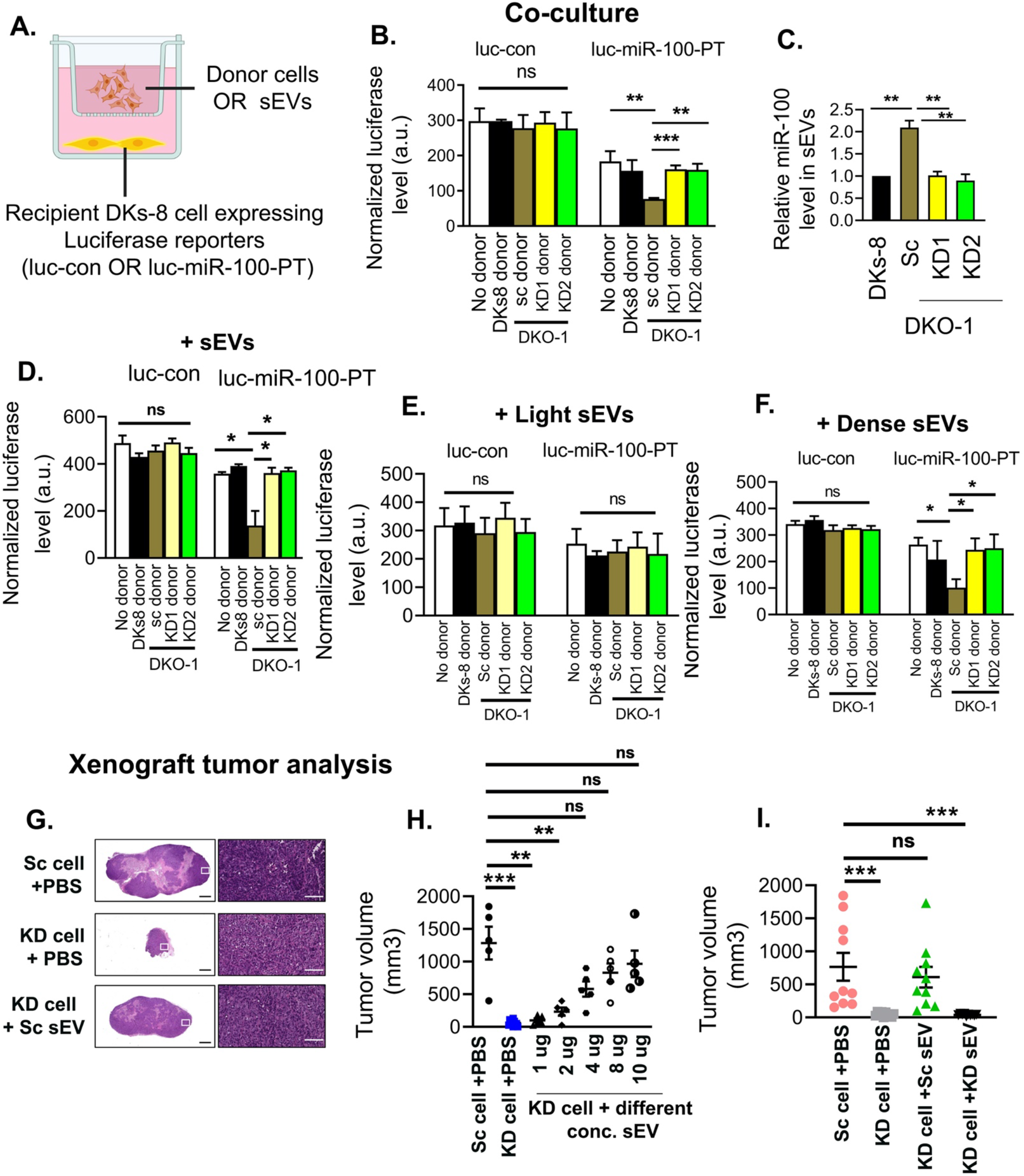
VAP-A controls miR-100 transfer and tumorigenic functions of EVs. (A) Illustration of co-culture setup. Control (luc-control: luciferase with scrambled sites in 3’ UTR) or miR-100 expressing luciferase reporters (luc-miR-100-PT: luciferase with miR-100 sites in 3’ UTR) (Cha et al., 2015) were expressed in recipient DKs-8 cells that were plated in the bottom of a Transwell plate. Different donor cells (DKs-8, or control or VAP-A KD DKO-1 cells) were cultured in Transwell inserts. Alternatively, sometimes purified EVs were added instead of donor cells. (B) Graph shows luciferase expression levels, normalized by co-expressed beta galactosidase, after lysis of recipient control or miR-100 reporter-expressing DKs-8 cells that were cocultured with the indicated donor cells. Data from three independent experiments. (C) Relative miR-100 levels in small EVs isolated from different donor cells, quantified by qRT-PCR. Data from three independent experiments. (D-F) Relative luciferase expression in recipient DKs-8 cells after addition of small EVs purified from donor cells, as indicated. (D), small EVs purified by cushion density gradient. (E and F) Light EVs or Dense EVs purified as in (Kowal et al., 2016). Luciferase data from three independent experiments with three technical replicates each time. (G-I) Control (Sc) and VAP-A KD2 (KD) DKO-1 cells were mixed with PBS (+PBS) or small EVs (+sEV) and injected subcutaneously in nude mice and allowed to grow for 3 weeks, with injection of PBS or EVs twice a week. (G) Representative images of Haematoxylin and Eosin stained sections of tumors. (H) Tumor volume after injecting PBS or different concentrations of small EVs purified from control (Sc) DKO-1 cells. (I) Tumor volume for control and KD tumors injected with PBS, or 10 μg control or KD sEVs, as indicated. Each condition from ten animals (I). Some data points in Figure H and I are in common. *p<0.05, **p<0.01, ***p<0.001. ns, not significant.

Our EV fractionation analysis in Fig 3 showed that the dense subpopulation of small EVs is enriched in RNA, including miR-100, and is regulated by VAP-A. To further validate that finding, we added light or dense small EVs purified from control or VAP-A KD DKO-1 cells, or from DKs-8 cells. Indeed, only the dense small EVs purified from control DKO-1 cells reduced luciferase expression in miR-100-PT-luciferase-expressing recipient cells (Figures 4E and F).

Previous reports showed that DKO-1 cells are tumorigenic when grafted into mice (Shirasawa et al., 1993). To test whether VAP-A-mediated EV production promotes tumor growth, we injected control and VAP-A KD DKO-1 cells into the flanks of nude mice and allowed tumors to grow for 21 days. Compared with control tumors, VAP-A KD tumors were much smaller or absent at the time of harvest (Figures 4G and 4H). To determine whether the defect in VAP-A KD growth was due to alterations in EV secretion, we performed a reconstitution experiment in which purified small EVs were mixed with VAP-A KD cells. Indeed, purified small EVs from control DKO-1 cells rescued the growth of KD tumors in a concentration dependent manner (Figure 4H). However, an equal amount of the highest concentration (10 μg) of EVs purified from VAP-A KD cells was not able to rescue VAP-A KD tumor growth (Figure 4I). These data suggest that VAP-A controls a specific subpopulation of EVs that promotes DKO-1 tumor growth.

### VAP-A controls the lipid content of EVs

VAP-A is known to promote efflux of lipids from the ER to diverse organelles by binding to lipid transporters, including oxysterol binding proteins (OSBPs) and ceramide transporters (Hanada et al., 2003; Mesmin et al., 2013; Perry and Ridgway, 2006). As ceramides, cholesterol and other lipids are thought to be involved in biogenesis of EVs, we hypothesized that VAP-A-mediated lipid transfer may be a critical component of the mechanism by which VAP-A promotes biogenesis of RNA-containing EVs. Analysis of free and total cholesterol levels in EVs revealed a significant reduction in both of those analytes in KD small EVs compared to control EVs (Figures 5A and B). By contrast, free and total cholesterol levels in KD cells are unchanged (Figures 5C and D). We also carried out an untargeted discovery lipidomics analysis of control and VAP-A KD small EVs, large EVs, and cells (Supplementary File Table 4). We found a variety of lipids predicted to be altered in KD EVs and cells. Notably, compared to controls, multiple ceramide species were decreased in both small and large KD EVs, with little or no change in KD cells (Supplementary File Table 4). We validated these findings for the most abundant ceramide species using targeted mass spectrometry with calibrated lipid standards. We found that C16.0, C22.0, and C24.1 ceramide are significantly reduced in VAP-A KD small and large EVs but unchanged in cells (Figures 5E-G).

**Figure 5:**
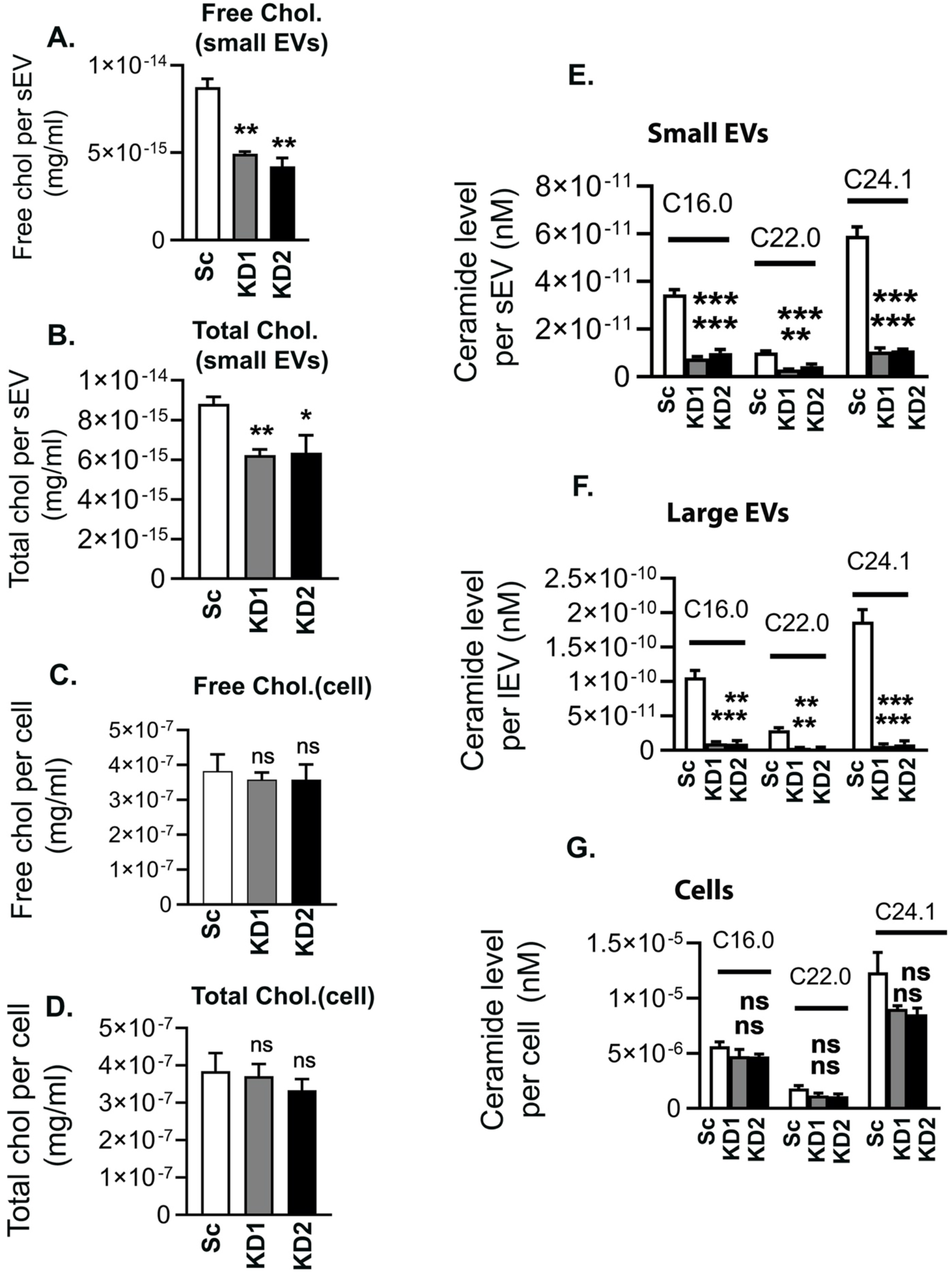
Cholesterol and ceramide levels are reduced in VAP-A KD EVs. (A and B) Free and total cholesterol level/EV for small EVs purified from control (Sc) and VAP-A KD DKs-8 cells. Data from three independent experiments. (C and D) Free and total cholesterol level/cell for control and KD DKs-8 cells. Data from three independent experiments. (E-G) Ceramide levels are reduced in VAP-A KD small and large EVs, but not in cells. Equal numbers of control and VAP-A KD small or large EVs, or cells were taken for ceramide (C16.0, C22.0, C24.1) measurements by targeted mass spectrometry. Data from three biological replicates. Graphs were plotted as mean ±S.E.M. *p<0.05, **p<0.01, ***p<0.001. ns, not significant.

### VAP-A binding partners CERT and ORP1L are critical for biogenesis of RNA-containing EVs

Since VAP-A interacts with CERT/STARD11 (Hanada et al., 2003) and VAP-A KD EVs have reduced ceramide levels, we tested whether CERT affects the RNA content of EVs by KD of CERT in DKs-8 cells. Similar to VAP-A KD, there is a significant effect of CERT-KD on the size and number of EVs released from cells (Figures 6A, 6B, S6A, and S6B). There is also a significant effect of CERT-KD on total RNA contents in small and large EVs (Figure S6C). Consistent with an important role for lipid transfer in the biogenesis of RNA-containing EVs, CERT-KD led to large reductions in the levels of candidate miRNAs in EVs but either no change or an increase in the levels of those same miRNAs in cells (Figures 6C-E).

**Figure 6:**
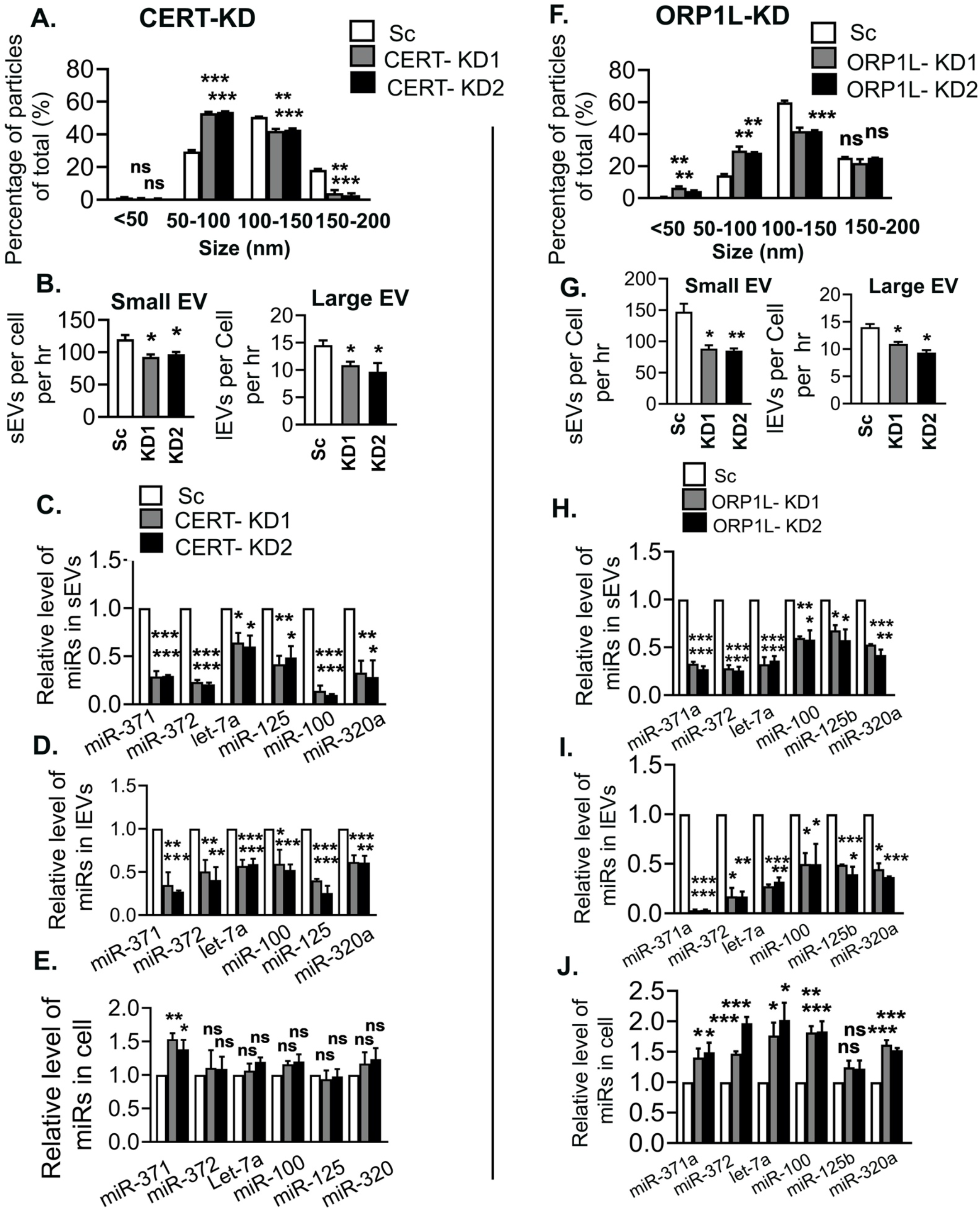
CERT and ORP1L control the biogenesis of miRNA-containing EVs. (A) NTA analysis of size distribution of small EVs purified from control (Sc) and CERT-KD (KD1, KD2) DKs-8 cells. Data from three independent experiments. (B) Small and large EV release rates from control and CERT-KD cells calculated from three independent NTA datasets. (C-E) qRT PCR analysis of miRNA levels in control and CERT-KD small and large EVs, and their parental cells, normalized to U6 snRNA. Data from three independent experiments. (F) NTA analysis of size distribution of small EVs purified from control (Sc) and ORP1L-KD (KD1, KD2) DKs-8 cells. Data from three independent experiments. (G) Small and large EV release rates from control and ORP1L-KD cells calculated from three independent NTA datasets. (H-J) qRT PCR analysis of miRNA levels in control and ORP1L-KD small and large EVs, and their parental cells, normalized to U6 snRNA. Data from three independent experiments. Data were plotted as mean +/- S.E.M. *p<0.05, **p<0.01, ***p<0.001. ns, not significant.

Since cholesterol may also affect the cargo content of EVs (Pfrieger and Vitale, 2018; Record et al., 2018) and is present at reduced levels in VAP-A KD sEVs, we tested whether KD of the endosomal VAP-A-binding cholesterol transporter ORP1L affects the RNA content of EVs. Similar to the results with CERT-KD, there was a reduction in the number, size, and total RNA content of ORP1L-KD small and large EVs (Figures 6F, 6G, and S6D-F). We also detected a large reduction of specific miRNAs in small and large EVs purified from ORP1L-KD cells, with either no change or an increase in cellular miRNA levels (Figures 6H-J). These data suggest that lipid transport from the ER to endosomes and the plasma membrane are critical for biogenesis and cargo content of RNA-containing EVs.

## Discussion

In this study, we examined the role of ER MCS in promoting biogenesis of RNA-containing EVs. Through both KD and overexpression experiments, we found that the ER MCS linker protein VAP-A promotes formation of RNA-containing small and large EVs. Small RNA-Seq analysis indicated that multiple classes of small non-coding RNAs are reduced in EVs purified from VAP-A KD cells. Likewise, lipid analysis of VAP-A KD EVs revealed a decrease in ceramides and other lipids, suggesting a potential mechanism for altered biogenesis of RNA-containing EVs. Indeed, KD of the VAP-A-binding lipid transporters ORP1L and CERT reduced the number of EVs as well as the relative RNA content. Biochemical subfractionation of EVs from control and VAP-A KD cells revealed that a previously characterized “dense” EV population contains the majority of RNA and is specifically downregulated in the KD EV population. This population appears to be important for miRNA transfer to recipient cells and tumor growth as EVs purified from VAP-A KD cells were not able to support those functions. Overall, we have identified a distinct subset of EVs that carries small noncoding RNAs and is formed at MCS by the action of VAP-A and VAP-A-binding lipid transporters.

Recent studies have shown that extracellular vesicles are released from cells as a heterogeneous population containing diverse protein cargoes (Kowal et al., 2016; Zhang et al., 2018). Leveraging a recently published method that subfractionates small EVs into light and dense populations (Kowal et al., 2016), we demonstrated that the dense population contains the minority of the small EVs (~10%) but is greatly enriched in RNA (~9-fold/EV) compared to light EVs. These data suggest that RNA-containing EVs are relatively rare in a general EV population, which may explain why previous calculations of RNA copies/EV are so low (Chevillet et al., 2014) despite their ability to transfer functional RNA to recipient cells (Abels et al., 2019; Chen et al., 2019; Ghamloush et al., 2019; Lucero et al., 2020; Shen et al., 2019; Ying et al., 2017). This subset of EVs was dependent on VAP-A expression in cells, as only the dense EV population was diminished in number, size, and RNA content with VAP-A KD. Our data further suggest that biogenesis of this EV population can be boosted and potentially regulated, since VAP-A overexpression greatly increased the number, size, and RNA content of EVs released from cells.

While RBPs are known to be important for the transport of RNAs into EVs (Leidal et al., 2020; Lin et al., 2019; Santangelo et al., 2016; Shurtleff et al., 2016; Villarroya-Beltri et al., 2013; Zietzer et al., 2020), it has been unclear how the RBP-RNA complexes are recruited to membranes for incorporation into EVs. Recent work has shown that two membraneless organelles that are comprised of RBP-RNA complexes - processing bodies and stress granules - form contacts with the ER (Lee et al., 2020). Furthermore, both biogenesis and fission of these organelles was shown to occur at these ER contact sites. Consistent with our localization of Ago2 and let-7a to ER-endosome contacts (Fig 1), one possibility is that RNA-containing membraneless organelles contact the ER at sites of EV biogenesis. Given our data that formation of RNA- and RBP-containing EVs depends on ER MCS proteins, one possibility is that fission or other dynamics of RNA-containing organelles at ER MCS may be coupled to biogenesis of RNA-RBP-containing EVs.

A major function of VAP-A at ER MCS is to promote lipid transport from one organelle to another. Indeed, we found that EVs purified from VAP-A KD cells had reductions in cholesterol, ceramide, and other lipids. In addition, we found that the ceramide and cholesterol transporters CERT and ORP1L were critical for biogenesis of RNA-containing EVs. Ceramide is known to be important for biogenesis of exosomes through induction of membrane curvature (Trajkovic et al., 2008). The role of cholesterol is less clear, although it is known to participate in the creation of ordered lipid microdomains that may recruit specific cargoes (Pfrieger and Vitale, 2018; Record et al., 2018). Thus, these two lipids may collaborate to couple cargo recruitment and EV biogenesis. Consistent with a model in which these processes occur at ER MCS, a recent paper showed that isolated ER membranes form contacts with other organelles at ordered lipid domains (King et al., 2020).

Lipids may also play a role in the binding of RNA-RBP complexes to ER MCS EV biogenesis sites. RNAs with a specific secondary structures bind to liquid-ordered domains in sphingomyelin–cholesterol–phosphatidylcholine vesicles (Janas et al., 2006) that resemble liquid-ordered lipid domains of exosomal membranes (Carayon et al., 2011). Interestingly, a specific exosome-sorting miRNA motif (EXO motif) reported earlier (Villarroya-Beltri et al., 2013) is present in the previously reported raft binding sequence (Janas et al., 2015; Janas et al., 2006). In addition, hnRNPA2B1 that is involved in sorting EXO-motif-containing miRNAs into exosomes colocalizes with ceramide-rich membrane regions (Villarroya-Beltri et al., 2013). Notably, we found that hnRNPA2B1 is one of the RBPs that depends on VAP-A for sorting into EVs. Once localized to ER MCS, it may be that additional molecular interactions facilitate incorporation of RNA-RBP complexes into EVs, such as the interaction with the ESCRT-0 protein Hrs that was recently shown to occur for FMR1 (Wozniak et al., 2020).

Our RNA sequencing data suggest that not all EV-associated RNAs are regulated by VAP-A mediated lipid transport. Indeed, U6, which is a small nuclear RNA commonly used to normalize miRNA levels in EVs (Cha et al., 2015; McKenzie et al., 2016) was not altered with VAP-A, CERT-, or ORP1L-KD. We do not believe it is a contaminant, because it was predominantly present inside of EVs, based on RNase sensitivity tests. Thus, it seems likely that there may be additional mechanisms that incorporate distinct RNAs into both small and large EVs. Nonetheless, biogenesis of RNA-containing EVs at ER MCS appears to be a major mechanism that controls specific sorting of miRNAs and a number of small noncoding RNAs. A recent study identified the autophagy protein LC3 to be involved in cargo loading into EVs, including RBPs hnRNPK and SAFB, and snoRNAs (Leidal et al., 2020). Whether LC3 is recruited to ER MCS to promote EV biogenesis or represents an alternative mechanism is an interesting topic for the future.

To test whether the subset of EVs that is controlled by VAP-A has functional relevance, we carried out several experiments. In miRNA transfer experiments, we found that VAP-A expression in donor DKO-1 colon cancer cells was critical for functional transfer of miR-100 to recipient DKs-8 colon cancer cells in both a co-culture setting and by direct addition of purified small EVs. Furthermore, dense but not light EV subfractions also mediated functional transfer of miR-100 to DKs-8 cells. We also carried out xenograft tumor experiments, in which we observed that VAP-A KD DKO-1 colon cancer cells had a defect in tumor growth. The rescue of that tumor growth defect by the addition of control but not VAP-A KD EVs indicates that the subpopulation of EVs controlled by VAP-A has important functional properties for tumor survival. Furthermore, it suggests that EV biogenesis at ER MCS is an important function of VAP-A in cancer cells.

In summary, we identified a novel biogenesis mechanism for RNA-containing small and large EVs that takes place at ER MCS. Our findings identify a new function for ER MCS, elucidate a poorly understood area of RNA and EV biology, and suggest pathways that could be leveraged for production of RNA-containing therapeutic EVs.

## ACKNOWLEDGMENTS

Thanks to the P01 group and Weaver laboratory for feedback and to Wade Calcutt for help with lipidomics. Funding was provided by NIH grants U19CA179514 and P01CA229123. Core facility usage (electron microscopy (Vanderbilt Cell Imaging Shared Resource), lipid mass spectrometry (Vanderbilt Mass Spectrometry Core) was supported in part by vouchers from Vanderbilt CTSA grant UL1 RR024975 and UL1 TR002243 and by NIH support of those facilities.

## AUTHOR CONTRIBUTIONS

BB designed and performed the majority of the experiments, carried out data analysis, and wrote the manuscript. JP performed bioinformatics analyses and edited the manuscript, EK provided expert advice, performed electron microscopy, and edited the manuscript, RA performed lipid analyses and edited the manuscript, KV provided expert advice and edited the manuscript, JGP provided reagents, expert advice, and edited the manuscript, QL provided expert advice and edited the manuscript, AMW aided in the experimental design, provided expert advice and co-wrote the manuscript.

## STAR*METHODS

### Key Resources Table

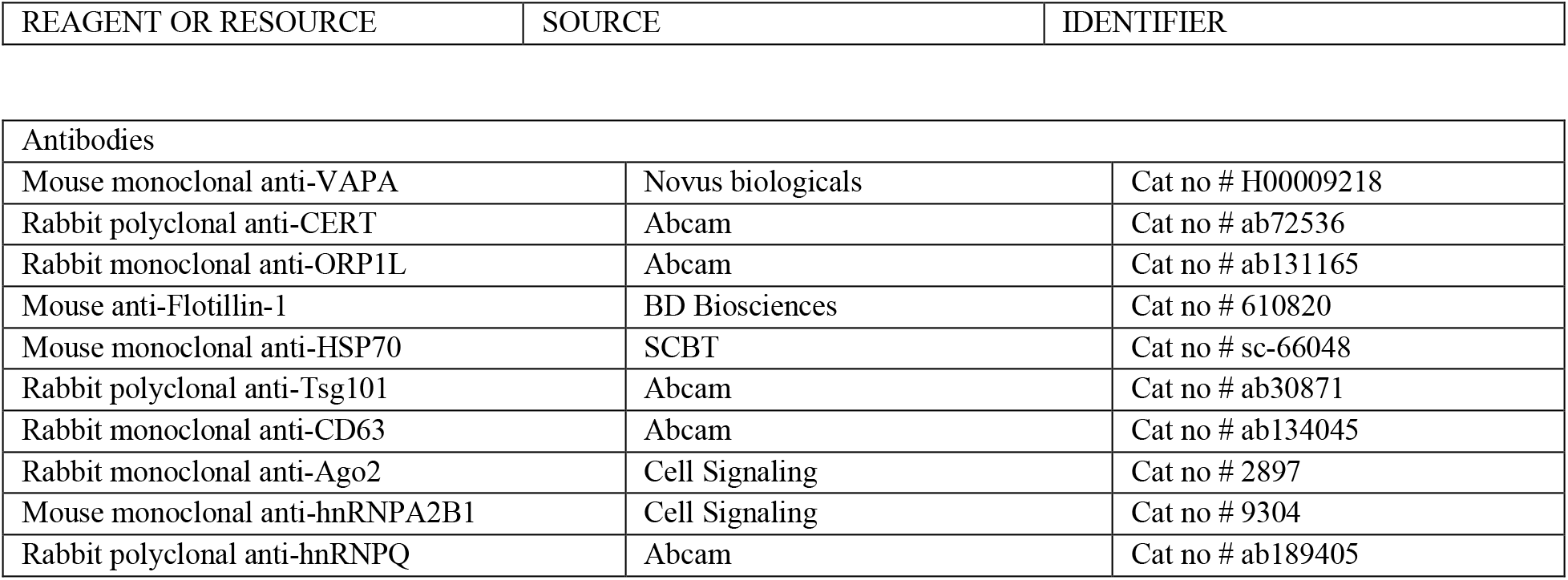
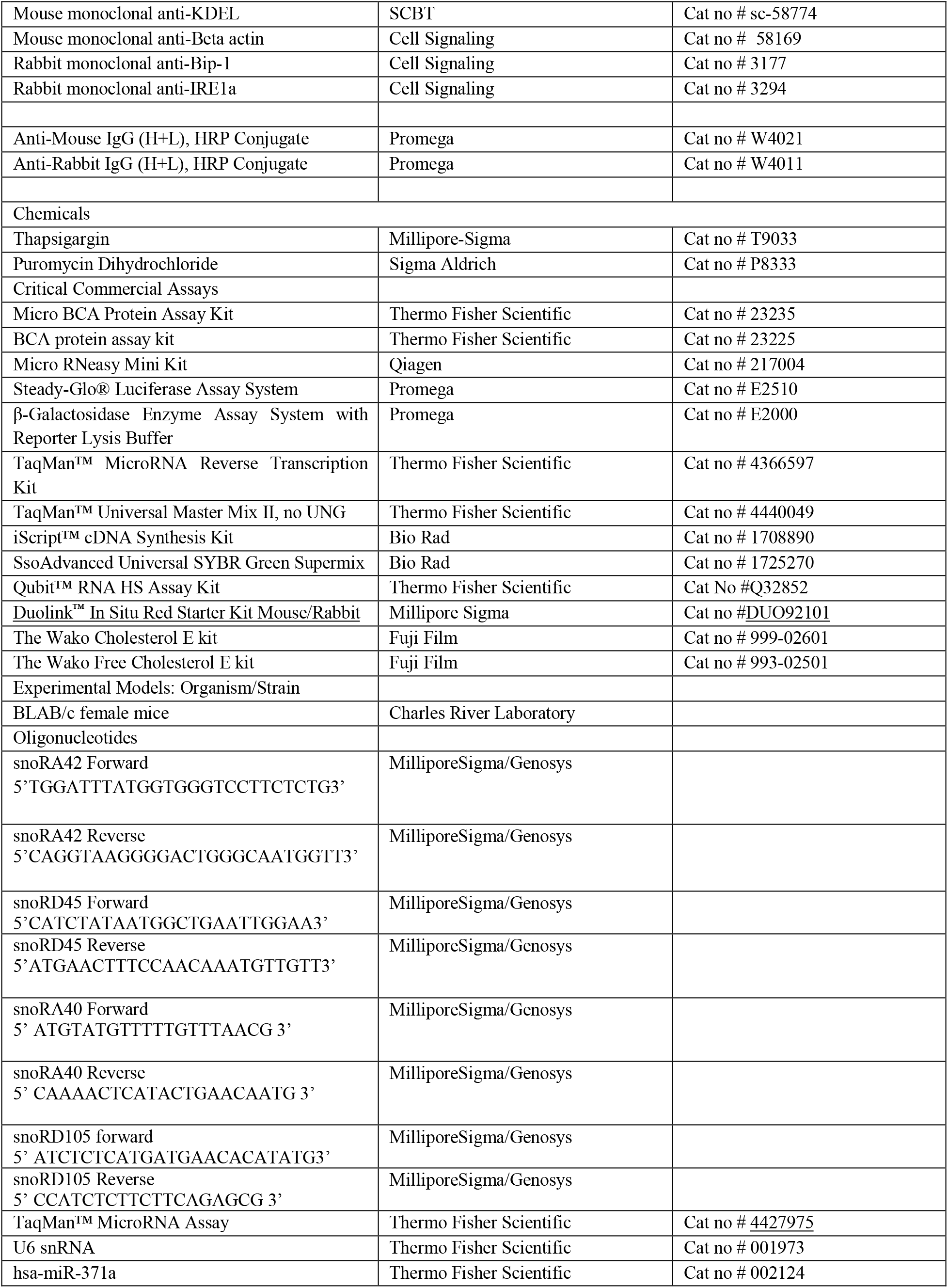
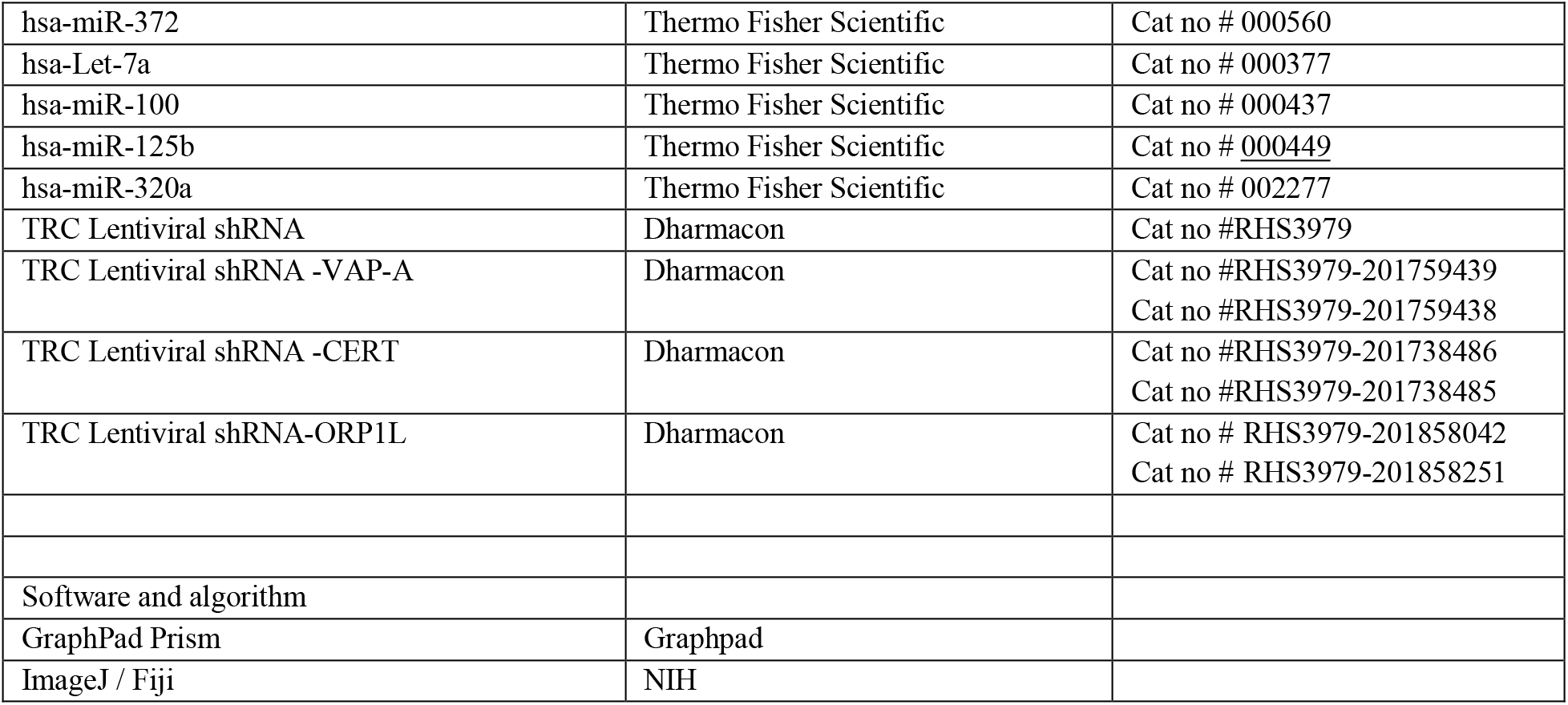

### RESOURCE AVAILABILITY

Lead contact and materials availability

Further information and requests for reagent and resources should be addressed to and will be met by the Lead Contact, Alissa Weaver. All unique/stable reagents generated in this study are available from the Lead Contact with a completed Materials Transfer Agreement.

### EXPERIMENTAL MODEL AND SUBJECT DETAILS

#### Cell Lines

WT DKs-8 and Mut DKO-1 were cultured in DMEM (Corning) supplemented with 10% Fetal bovine serum (FBS), non-essential amino acids (Sigma), and L-glutamine. HEK293FT cells were cultured in DMEM supplemented with 10% FBS and sodium pyruvate as per the manufacturer’s instructions.

#### Animal subjects

7-12 weeks old female athymic nude mice were purchased from Charles River Laboratory and kept in a pathogen-free facility approved by the American Association for the Accreditation of Laboratory Animal Care that met all current regulations and standards of the U.S. Department of Agriculture, U.S. Department of Health and Human Services, and the National Institutes of Health. Mice were fed irradiated standard mouse chow (LabDiet) and autoclaved, reverse osmosis treated water.

#### Non-orthotopic nude mouse model for tumor cell xenograft

Subconfluent cultures were harvested by trypsinization and washed with PBS. Subcutaneous tumors were established by injecting cells (7 × 10^6^ control or VAP-A-KD DKO-1 cells) suspended in 150 μL of serum-free DMEM into the flanks of nude mice as described (Ref. if any). In some cases small EVs or PBS was mixed with the cells before implantation and small EVs or PBS was injected twice a week until tumor harvest. Mice were examined twice a week for tumor size and weight loss. Subcutaneous tumor size was measured with micro calipers. Tumor volume was calculated as (A) X (B^2^) X 0.52 where A is the longest dimension of the tumor and B is the dimension of the tumor perpendicular to A. Mice were sacrificed after 3 weeks and tumors were fixed, sectioned, and stained with haematoxylin and eosin (H&E). Imaging of H&E stained tumor sections was performed using an Aperio Versa 200 scanner (Leica) in the Vanderbilt Digital Histology Shared Resource.

## METHOD DETAILS

### Extracellular vesicle isolation and nanoparticle tracking analysis

For cushion density gradient method, DKs-8 and DKO-1 cells were cultured at 80% confluence in serum-free DMEM. After 48 hours, the conditioned medium was collected from the cells and the EVs were isolated via serial centrifugation. Floating live cells, dead cell debris, and large EVs were respectively collected from the conditioned medium by centrifugation at 300 × g for 10 min, 2,000 × g for 25 min, and 10,000 × g Ti45 rotor, Beckman Coulter)) for 30 min. The supernatant was then overlaid onto a 2 ml 60% iodixanol cushion and centrifuged at 100,000 × g (SW32 rotor, Beckman Coulter) for 18h. The bottom 3 ml, including the 1 ml of collected EVs + 2 ml iodixanol (40% iodixanol final concentration) were transferred to the bottom of another tube and then 20%, 10% and 5% iodixanol were layered successively on top. These iodixanol dilutions were prepared by diluting OptiPrep (60% aqueous iodixanol) with 0.25 M sucrose/10 mM Tris, pH 7.5. After an 18-hour centrifugation step at 100,000 × g, 12 density gradient fractions were collected, diluted in PBS and centrifuged at 100,000 × g for 3 hours. To quantitate the size and concentration of EVs, nanoparticle tracking analysis (NTA) was performed using a Particle Metrix ZetaView PMX 110.

For “light” and “dense” small EV purification, pellets obtained by ultracentrifugation (100,000 x g in Ti45 Beckman Coulter rotor for 70 min. at 4 ° C) were washed and resuspended in 1.25 ml buffer [0.25 M sucrose, 10 mM Tris pH 8.0, 1 mM EDTA (pH 7.4)] and transferred to a SW55Ti rotor tube (Beckman Coulter), and mixed with 60% (wt/vol) stock solution of iodixanol (1:1). Next, 1.1 ml 20% (wt/vol) iodixanol and 1 ml of 10% (wt/vol) iodixanol successively layered on top of the vesicles suspension and tubes were centrifuged for 1h at 4 °C at 350,000 x g in SW55Ti rotor; Ten fractions of 460 ul were collected from the top. Fractions were diluted and washed in PBS for 1h at 100,000 x g in a TLA 110 rotor (Beckman). Fractions were resuspended in 35 ul of PBS. This method was from a previously published report (Kowal et al., 2016).

### Fluorescence microscopy

Cells on coverslips coated with poly-D-Lysine (100 μg/ml) were fixed with 4% paraformaldehyde in PBS then permeabilized with 0.5% Triton X-100 in PBS. Cells were stained for proximity ligation assay (PLA) using a Duolink® kit according to the manufacturer’s protocol (DUO92101, Millipore Sigma). Briefly, cells were blocked by Duolink® Blocking Solution for 60 minutes in a 37° C humid chamber. Primary antibodies were diluted (KDEL 1:100 and CD63 1:100) in the Duolink® Antibody Diluent and incubated overnight at 4° C in a humid chamber. After washing, the cover slips were incubated with PLA probes for 1h in a 37° C humid chamber. The ligation reaction was performed for 30 minutes at 37° C followed by washing and amplification at 37 degree Celsius for 100 minutes. Cover slips were washed and mounted with <u>Duolink® In Situ Mounting Medium with DAPI</u> provided in the kit. Single confocal images were acquired with a Nikon A1R-HD25 confocal microscope (NIS-Elements) equipped with an Apo 60x/1.49 NA oil immersion lens. Images were analyzed for colocalization of PLA dots (generated by KDEL and CD63 proximity) with GFP-Ago2 or Cy3-let7 fluorescence signals surrounding cells were segmented from the background by thresholding and measured for colocalization by Fiji (Analyze tab/Colocalization/Colocalization Threshold) and Pearson’s coefficient was taken for plot. For florescent dots quantification, images were segmented from the background by thresholding and calculate particle number (Analyze/analyze particles) per cell by Fiji.

### Transmission electron microscopy (TEM)

For negative staining of purified EVs, Formvar carbon film-coated grids (FCF-200-Cu; Electron Microscopy Sciences) were washed in double distilled water and then washed by 100% ethanol. For each step, excess liquid was removed by wicking with filter paper. 10-μl samples were added to grids overnight at 4 ° C. Grids were then incubated with 2% phosphotungstic acid, pH 6.1 for 30 s and followed by blotting by touching filter paper to the side of the grid.

For TEM of the cells, samples were fixed in 1.3% glutaraldehyde in 0.1 M cacodylate for 1 hour at room temperature followed by 24 hours at 4 ° C. Samples were post-fixed in 2% OsO4, 1% potassium ferrocyanide for 1 hour and en bloc stained with 0.5% uranyl acetate. Samples were dehydrated with a graded ethanol series, infiltrated with Epon-812 (Electron Microscopy Sciences) using propylene oxide as a transition solvent, and polymerized at 60 ° C for 48 hours. Samples were sectioned at a nominal thickness of 70 nm and stained with 2% uranyl acetate and lead citrate.

All TEM samples were imaged using a Tecnai T12 operating at 100 kV with an AMT CCD camera using AMT imaging software. Analysis of the TEM data was performed in FIJI.

### Western blot analysis

The protein concentrations of total cell lysates were determined utilizing Pierce BCA Assay (Cat. 23225, Thermo Fisher). The protein concentrations of the EVs were determined utilizing Pierce Micro BCA Assay (Cat. 23235, Thermo Fisher). For Western blots, 15 μg of TCLs, small EVs, large EVS or an equal volume of resuspended vesicles from density gradient fractions (for control markers blots) were boiled in SDS-Page sample buffer for 5 min and loaded on 10-well or 15-well 8% or 10% polyacrylamide gels. Proteins were transferred to nitrocellulose membranes for 1 h at 100 volts or 25 volts for overnight at 4°C. Membranes were blocked in 3% BSA diluted in Trisbuffered saline with 0.5% Tween 20 (TBST) for 4h at room temperature. Primary antibodies were diluted in 3% BSA -TBST (Ago2, 1:1000; hnRNPA2/B1,1:1000; Hsp70, 1:1000; CD63, 1:1000; Flotillin, 1:1000; TSG101, 1:1000; GM130, 1:2000; and beta actin, 1:10000) and incubated overnight at 4°C. Membranes were washed 3 times for 15 min in TBST and subsequently incubated with species-specific HRP-conjugated secondary antibodies (1:10000; Promega) in 3% BSA-TBST for 1h at room temp. All membranes were washed 3 times for 15 min in TBST and incubated with an enhanced chemiluminescence (ECL) reagent (Thermo Scientific) for 1 min before being exposed to film or using a ChemiDoc Imager (BioRad) or Amersham 680 imager (GE). Multiple exposures were taken for each blot to have the complete dynamic range for densitometry measurements. The densitometry measurements for the protein bands were done using the Analysis Gels feature of ImageJ (NIH).

### Dot blot analysis

Dot blotting of EV preparations was performed as described previously (Lai et al., 2015). Different concentrations of sEVs or lEVs were collected from conditioned media of DKs-8 cells were dotted onto nitrocellulose membranes and allowed to dry at room temperature for 1 h. The membrane was - blocked with 3% BSA in Tris-buffered saline (TBS) in the absence or presence of 0.1% (v/v) Tween 20 (TBS-T) at room temperature for 1h, followed by incubation with anti-Ago2, anti-hnRNPA2/B1, anti-flotillin-1 or antiCD63 antibody in TBS or TBS-T overnight at 4 °C.

### RNA purification

Total RNA from cell, small and large EVs was purified using the miRNeasy kit (Qiagen Inc., Valencia, CA, USA) according to the manufacturer’s protocol. Final RNAs were eluted with two rounds of 35 ul of Nuclease free water extraction.

### miRNA library preparation and sequencing

Total RNA from each sample was used for small RNA library preparation using NEBNext. Small RNA Library Prep Set from Illumina (New England BioLabs Inc., Ipswich, MA, USA). Briefly, 3’ adapters were ligated to total input RNA followed by hybridization of multiplex single read (SR) reverse transcription (RT) primers and ligation of multiplex 5’ SR adapters. RT was performed using ProtoScript II RT for 1 hr at 50°C. Immediately after RT reactions, PCR amplification was performed for 15 cycles using LongAmp Taq 2× master mix. Illumina-indexed primers were added to uniquely barcode each sample. Post-PCR material was purified using QIAquick PCR purification kits (Qiagen Inc.). Post-PCR yield and concentration of the prepared libraries were assessed using Qubit 2.0 Fluorometer (Invitrogen, Carlsbad, California, CA, USA) and DNA 1000 chip on Agilent 2100 Bioanalyzer (Applied Biosystems, Carlsbad, CA, USA), respectively. Size selection of small RNA with a target size range of approximately 146–148 bp was performed using 3% dye free agarose gel cassettes on a Pippin Prep instrument (Sage Science Inc., Beverly, MA, USA). Post-size selection yield and concentration of libraries were assessed using Qubit 2.0 Fluorometer and DNA high-sensitivity chip on an Agilent 2100 Bioanalyzer, respectively. Accurate quantification for sequencing applications was performed using qPCR-based KAPA Biosystems Library Quantification kits (Kapa Biosystems, Inc., Woburn, MA, USA). Each library was diluted to a final concentration of 1.25 nM and pooled in equimolar ratios prior to clustering.

Single-end sequencing was performed to generate at least 15 million reads per sample on an Illumina HiSeq2500 v4 using a 50-cycle kit.

### Small RNA sequencing analysis

Small RNA-seq reads were trimmed using cutadapt v1.18 (https://github.com/marcelm/cutadapt). After trimming, reads longer than 15 nucleotides were retained. ncPRO-seq (version 1.5.1) (Chen et al., 2012) was used to map reads to the reference genome hg19 and quantitate small RNA. The miRBase v18, ACA_snoRNA and CD_snoRNA from Rfam v11.0, and tRNA from UCSC (hg19) were employed for reads annotation to miRNA, snoRNA, and tRNA. miRNA annotation was extended in both upstream and downstream regions by using miRNA_e_+2_+2. Principal component analysis was performed to assess the similarity between samples. DESeq2 (Love et al., 2014) was used to identify small RNAs differentially exported from Cell to EVs in VAP-A KD compared to SC. For identifying miRNAs that were upregulated or downregulated in KD EVs or cells, all KD samples were compared to control in one group whether from KD1 or KD2 cells. Small RNAs with fold change≥2 or ≤0.5 and a false discovery rate (FDR) ≤ 0.05 were considered to be significantly differentially expressed.

### qRT-PCR for miRNA

Total RNA was isolated from small EVs, large EVs, and cells using the miRNeasy kit (Qiagen Inc., Valencia, CA, USA), which isolates all small RNAs <200 nt, including miRNAs. Total RNA amount of sEVs, lEVs and Cells were measured by Nanodrop. Taqman small RNA assays (ThermoFisher Scientific) were performed for small EVs, large EVs and cellular RNAs according to the manufacturer’s protocol; U6 snRNA: 001973; hsa-let7a-5p: 000377; hsa-miR-100-5p:000437; hsa-miR320a: 002277; has-miR-371a: 002124; has-miR-372: 000560. Individual reverse transcription reactions were performed using 10 ng RNA from each sample per Taqman miRNA primer in a final reaction volume of 10 μl. After transcription, 0.34 ng (0.67 μl) cDNA was used as the template together with the corresponding Taqman miRNA probe for qPCR in a final reaction volume of 10 μl. Each Taqman miRNA qPCR was performed with technical triplicates on a Bio-Rad CFX96. C(t) values were averaged for each technical triplicate. U6 snRNA was used as a normalization control for each biological sample. To calculate fold changes (FC), the ΔΔC(t) method was used (Schmittgen and Livak, 2008). Briefly, ΔC(t) values were calculated for each biological sample, where ΔC(t) = C(t)miRNA - C(t)U6 snRNA. Relative fold changes were determined by Fold change = 2-ΔΔC(t), where ΔΔC(t) = ΔC(t)-ΔC(t)control. For ΔΔC(t) values < 0 (signifying a negative fold change), the negative reciprocal Fold Change formula was used (−1/(2-ΔΔC(t)). Statistical analyses were performed from three independent biological replicates

### RNase protection assay for EV samples

EV pellets resuspended in PBS were mixed with Triton-X-100 (TX-100) (final concentration 1%) and 10 Units RNase I (Thermo) in 100 μl for 30 min at 37 ° C. Enzyme was inactivated at 95 ° C for 10 min and 700 μl Trizol was added followed immediately by RNA extraction using the miRNeasy kit (Qiagen Inc., Valencia, CA, USA).

### Co-culture and Luciferase reporter assay

Recipient DKs-8 cells were plated in six-well plates at a density of ~2.5 × 10^5^ cells and cultured in DMEM supplemented with 10% FBS for 24 hr. The media was replaced with serum-free Opti-MEM and the cells were co-transfected with 1.5 μg of Luc-reporter plasmid and 1.5 μg β-gal plasmid DNA/well. Donor cells were plated in 0.4-μm pore Transwell filters (Corning, 3450, Corning, NY, USA) at ~2.5 × 10^5^ cells/well for 24 hr. The media from donor Transwells and recipient 6-well plates were removed and replaced with serum-free DMEM. Co-culture of donor and recipient cells was then conducted for 48 hr before recipient cells were harvested. In some cases, purified small EVs were added instead of co-culturing with donor cells (8X10^9^ per well for Fig 4D, 2X10^9^ per well for Fig 4E,F). The number of EVs to add was estimated by the EV/cell/hour secretion rate of parent DKO-1 cells x number of cells x number of hours of assay then refined in pilot experiments. Lysates were prepared in 1 × Reporter lysis buffer (Promega, E2510), and Luciferase assays were performed according to the manufacturer’s protocol (Promega, E2510). β-gal expression was simultaneously determined from the lysates according to the manufacturer’s protocol (Promega, E2000). Differences in transfection efficiency were accounted for by normalizing Luc expression to β-Gal expression (Luc/β-Gal). All assays were performed on three biological replicates, each with three technical replicates.

### Lipid isolation and cholesterol assay

Lipid contents from cell and EVs were extracted using the Bligh-Dyer method (Bligh and Dyer, 1959). Total lipids were isolated from cell or EVs with chloroform: methanol: saline (1:1:0.9) extraction. Lipids were extracted from organic phase (bottom) by evaporating chloroform under N2 gas followed by resuspension in chloroform containing 1% Triton-X-100 detergent and reevaporating of chloroform under N2 gas. Finally, the mixture was resuspended in saline. Lipids assays were done using colorimetric kits for total cholesterol (Wako; 999-02601) and free cholesterol (Wako; 993-02501) and normalized to small EV numbers or cell numbers. Cholesterol level was not normalized by protein levels since protein levels were also altered between control and VAP-A KD small and large EVs (data not shown).

### Lipid mass spectrometry

Untargeted Lipidomics. Discovery lipidomics data were acquired using a *Vanquish* ultrahigh performance liquid chromatography (UHPLC) system interfaced to a *Q Exactive HF* quadrupole/orbitrap mass spectrometer (Thermo Fisher Scientific). Samples were injected a total of four times. Two injections were made in positive ESI mode followed by two injections in negative mode. Chromatographic separation was performed with a reverse-phase Acquity BEH C18 column (1.7 mm, 2.1×150mm, Waters, Milford, MA) at a flow rate of 300 ml/min. Mobile phases were made up of 10 mM ammonium acetate in (A) H_2_O/CH_3_CN (1:1) and in (B) CH3CN/iPrOH (1:1). Gradient conditions were as follows: 0–1 min, B = 20 %; 1–8 min, B = 20–100 %; 8-10 min, B = 100 %; 10–10.5 min, B = 100–20 %; 10.5-15 min, B = 20%. The total chromatographic run time was 20 min; the sample injection volume was 10 mL. Mass spectra were acquired over a precursor ion scan range of *m/z* 100 to 1,200 at a resolving power of 30,000 using the following ESI source parameters: spray voltage 5 kV (3 kV in negative mode); capillary temperature 300°C; S-lens RF level 60 V; N2 sheath gas 40; N2 auxiliary gas 10; auxiliary gas temperature 100 °C. MS/MS spectra were acquired for the top-five most abundant precursor ions with an MS/MS AGC target of 1e5, a maximum MS/MS injection time of 100 ms, and a normalized collision energy of 30. Chromatographic alignment, peak picking, and statistical comparisons were performed using Compound Discoverer v. 3.0 (Thermo Fisher). All differential features (samples vs. controls) having a *P* value of <0.05 and a fold change of >1.5 were processed for molecular matches in the Chemspider, mzCloud, HMDB, and KEGG databases based on precursor ion exact masses (+/-5 ppm) and MS/MS fragmentation patterns. Lipid matches and identified metabolites were then filtered to exclude biologically irrelevant drugs and environmental contaminants, and the finalized list of putative identifications were mapped to relevant biological pathways using the Metabolika software module. Pooled QCs were injected to assess the performance of the LC and MS instruments at the beginning and at the end of each sequence.

#### Targeted Lipidomics

The triple quadrupole mass spectrometer was operated in positive ion mode. Quantitation was based on multiple reaction monitoring detection. The following optimized source parameters were used for the detection of analyte and internal standards. Ceramides: N2 15 sheath gas 60 psi; N2 auxiliary gas 5 psi; spray voltage 3.5 kV; HESI probe temperature 386 °C; capillary temperature 300 °C; declustering voltage 5 V. Calibration curves were constructed for ceramides C16:0, C22:0 and C24:1 by plotting peak area ratios (analyte / internal standard) against analyte concentrations for a series of ten calibrants, ranging in concentration from 10 ng/mL to 20 μg/mL. A weighting factor of 1/Ct 2 was applied in the linear least-squares regression analysis to maintain homogeneity of variance across the concentration range (% error ≤ 20% for at least four out of every five standards). Data acquisition and analysis were carried out using Xcalibur v.2.1.0, Vantage v.2.3.0 and LCQuan v.2.7.0 software (Thermo).

### Non-orthotopic nude mouse model for tumor cell xenograft

Subconfluent cultures were harvested by trypsinization and washed with PBS. Subcutaneous tumors were established by injecting cells (7×10^6^ control or VAP-A-KD DKO-1 cells) suspended in 150 μL of serum-free DMEM into the flanks of nude mice as described (Ref. if any). In some cases, small EVs (1×10^11^ to 10×10^11^ EVs) or PBS were mixed with the cells before implantation and small EVs or PBS were injected twice in a week until tumor harvest. The number of EVs to add was first estimated from the EV secretion rate x number of cells x hours before next injection then converted to protein, for ~4 μg. Pilot experiments then tested 1-10 μg protein concentrations (Fig 4H). Mice were examined twice a week for tumor size and weight loss. Subcutaneous tumor size was measured with micro calipers. Tumor volume was calculated as (A) X (B^2^) X 0.52 where A is the longest dimension of the tumor and B is the dimension of the tumor perpendicular to A. Mice were sacrificed after 3 weeks and tumors were fixed, sectioned, and stained with haematoxylin and eosin (H&E). Imaging of H&E stained tumor sections was performed using an Aperio Versa 200 scanner (Leica) in the Vanderbilt Digital Histology Shared Resource.

### Statistics

Experimental data were acquired from at least three independent experiments. Data plotted by bar graph were compared using student’s *t* test and plotted as mean and standard error of the mean using GraphPad Prism 7. Data plotted as box and whisker plots were checked for normality and analyzed by one way ANOVA with Bonferroni’s multiple comparison test (Fig 1E) or with Welch’s correction (does not assume equal standard deviation (Fig 1F)), or assessed for Pearson’s correlation (Fig S1A). Tumor data were compared by Mann-Whitney test.

---

**Figure S1.**
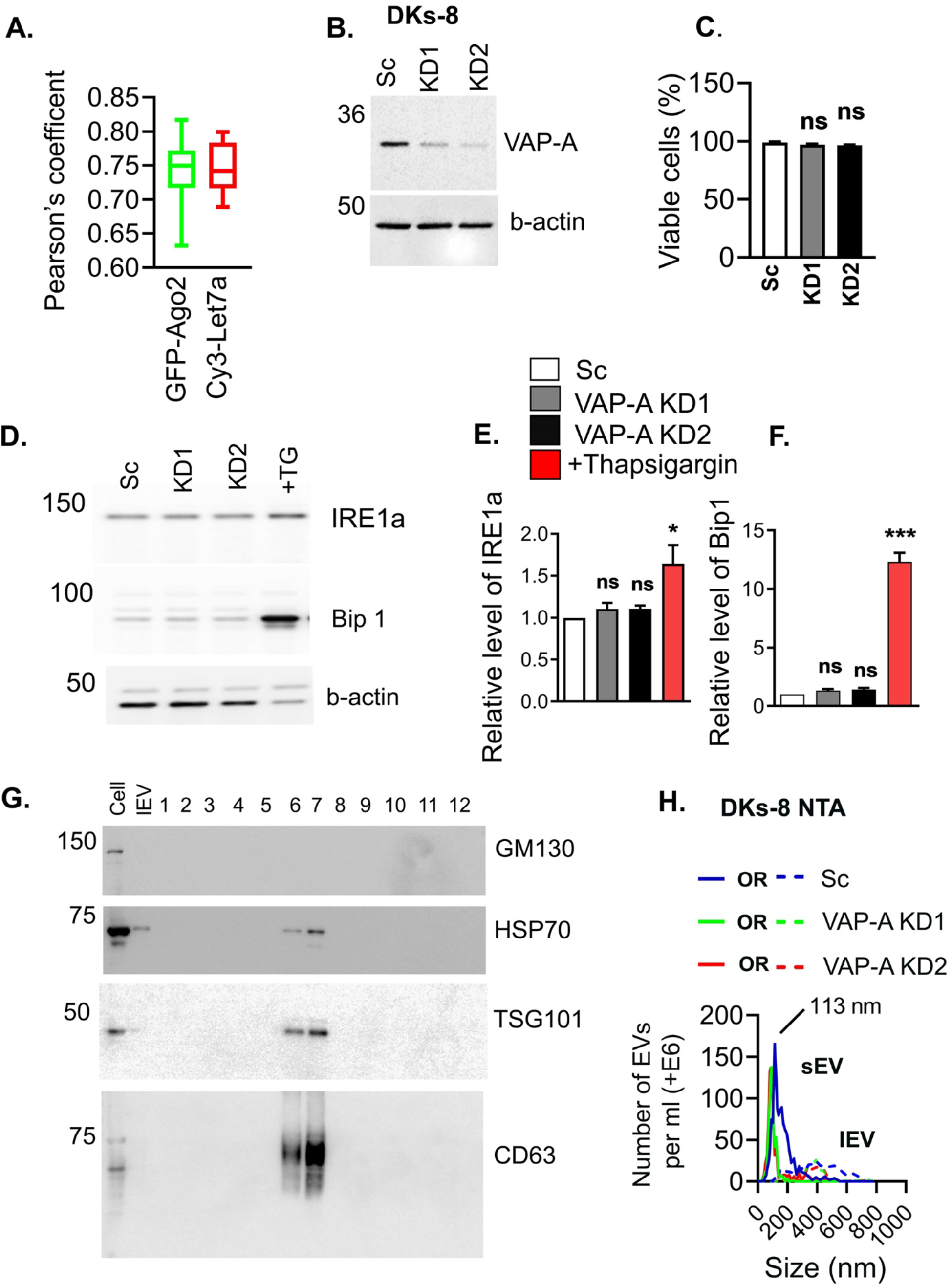
Characterization of EVs and ER stress upon VAP-A KD in DKs-8 cells. Related to Figure 1. (A) Pearson’s quantification of colocalization of ER MCS PLA with Ago2 or let-7a, related to images shown in Figures 1C and 1D. (B) Western blot analysis of VAP-A levels in control (Sc) and VAP-A KD DKs-8 cells. Beta actin serves as an endogenous control. (C) Percent viability of control (Sc), VAP-A KD (KD1and KD2) DKs-8 cells at the time of conditioned media collection. Data from three independent experiments. (D-F) Analysis of ER stress by Western blot. (D) Representative immunoblots show ER stress markers (Bip-1 and IRE1a) in control (Sc) and VAP-A KD DKs-8 cells. Thapsigargin (ER stress inducer; 10 μM, overnight) treatment of control cells serves as a positive control. Beta-actin serves as an endogenous control. Quantification of IRE1a (E) or Bip1 (F) was performed for three independent experiments. (G) Western blot analysis of positive EV markers (Hsp70, Tsg101 and CD63) and a negative EV marker (GM130) in DKs-8 cells, large EVs (lEV) and small EVs (fractions 6 and 7 of the density gradient). (H). Representative nanoparticle tracking analysis (NTA) traces of small (sEV) and large (lEV) EVs purified from control (Sc) and VAP-A KD (KD1 and KD2) DKs-8 cells. Data plotted as Mean ± S.E.M. *p<0.05, **p<0.01, ***p<0.001.

**Figure S2:**
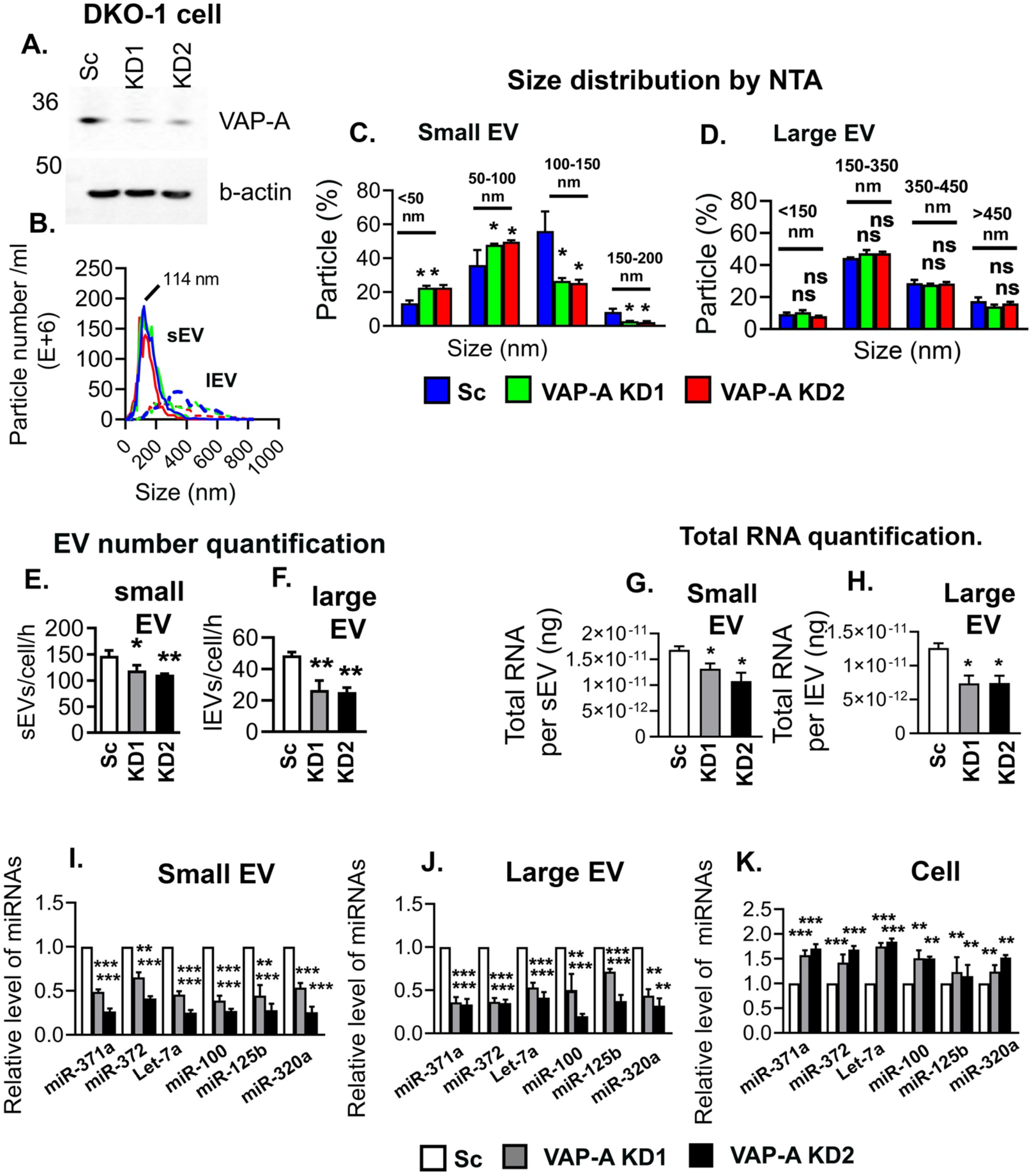
VAP-A promotes biogenesis of RNA-containing EVs in DKO-1 cells. Related to Figures 1 and 2. (A) Western blot of VAP-A in DKO-1 colon cancer cells shows KD of VAP-A. Beta actin serves as an endogenous control. (B) Representative nanoparticle traces for small and large EVs purified from control (Sc) and VAP-A KD DKO-1 cells. (C and D) Analysis of size distribution of small and large EVs measured by NTA from control (Sc) and VAP-A KD (KD1 and KD2) DKO-1 cells. Particles from three independent experiments were binned according to their size for plotting and statistical analysis. (E and F) EV secretion rates calculated from NTA data for EVs isolated from control (Sc) and VAP-A KD DKO-1 cell conditioned media. Data from three independent experiments. (G and H) Total RNA was extracted from a known number of purified EVs and measured by NanoDrop (A260). The concentration of RNA was plotted per small or large EV. Data from three independent experiments. (I-K) Graphs show relative level of specific miRNAs quantified by qRT-PCR in small and large EVs purified from control (Sc) and VAP-A KD DKO-1 cells and from their respective parental cells. Data from three independent experiments. Data were plotted as Mean ± S.E.M. p*<0.05, p**<0.01, p***<0.001.

**Figure S3.**
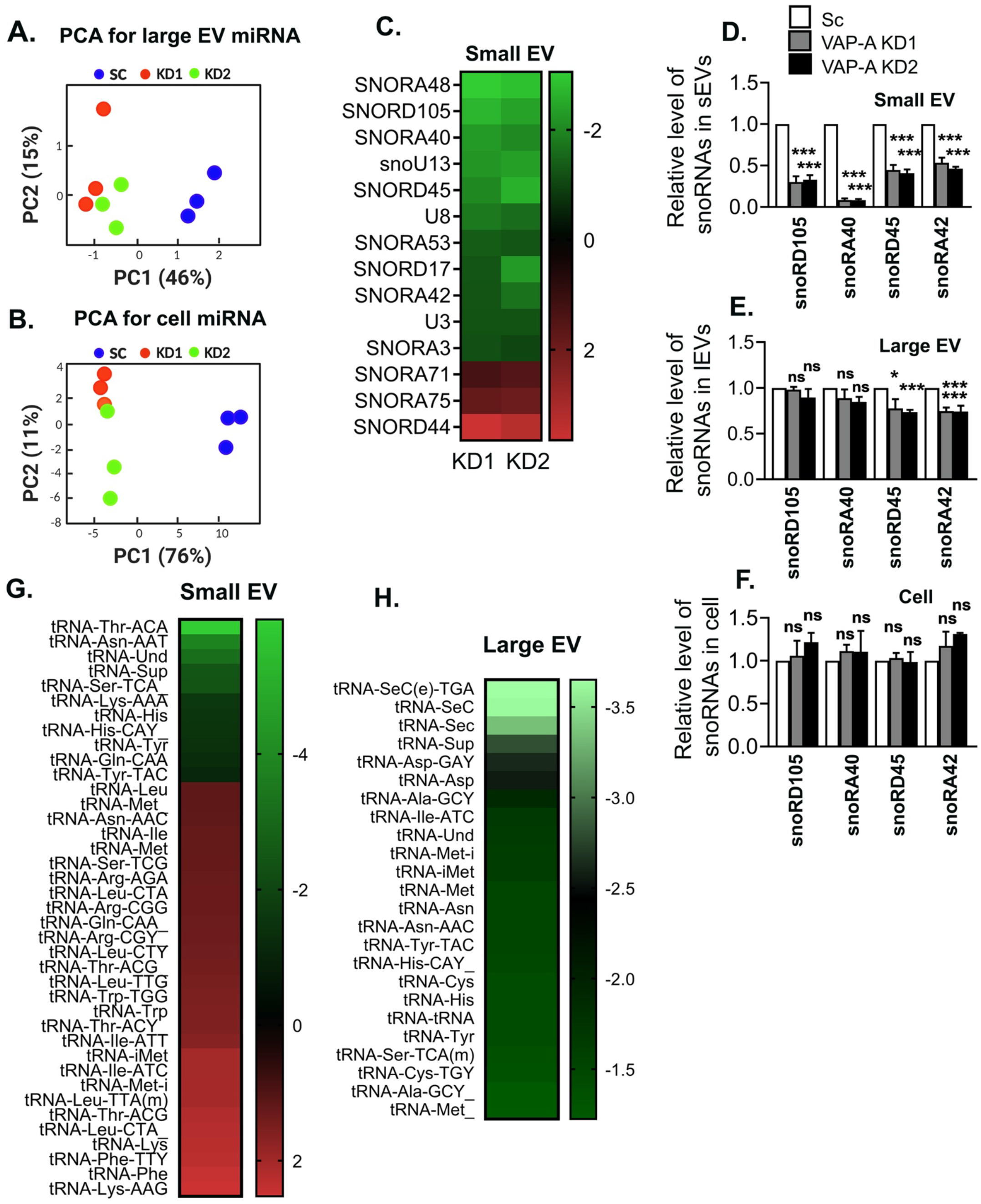
VAP-A regulates the levels of small RNAs in EVs. Related to Figure 2. (A and B) Principal component analysis (PCA) shows that VAP-A KD affects the miRNA composition of large EVs and cells. (C) Heatmap analysis of small RNA-Seq data shows altered snoRNAs in VAP-A KD small EVs purified from DKs-8 cell using criteria ≤ 0.5 or ≥ 2 fold change and FDR value<0.05. Green shows downregulated whereas red shows upregulated RNAs. Levels plotted as log 2-fold change. (D-F) Relative levels of specific snoRNAs quantified by qPCR and normalized to U6 snRNA in small EVs, large EVs and their parental control (Sc) or VAP-A KD DKs-8 cells, as indicated above. Data from three independent experiments. (G and H) Heatmap analyses of small RNA-Seq data show altered tRNA fragments in small and large EVs in VAP-A KD conditions, using criteria ≤ 0.5 or ≥ 2 fold change and FDR value<0.05. Levels plotted as log 2-fold change. Histograms are plotted as Mean ± S.E.M. *p<0.05, **p<0.01, ***p<0.001.

**Figure S4.**
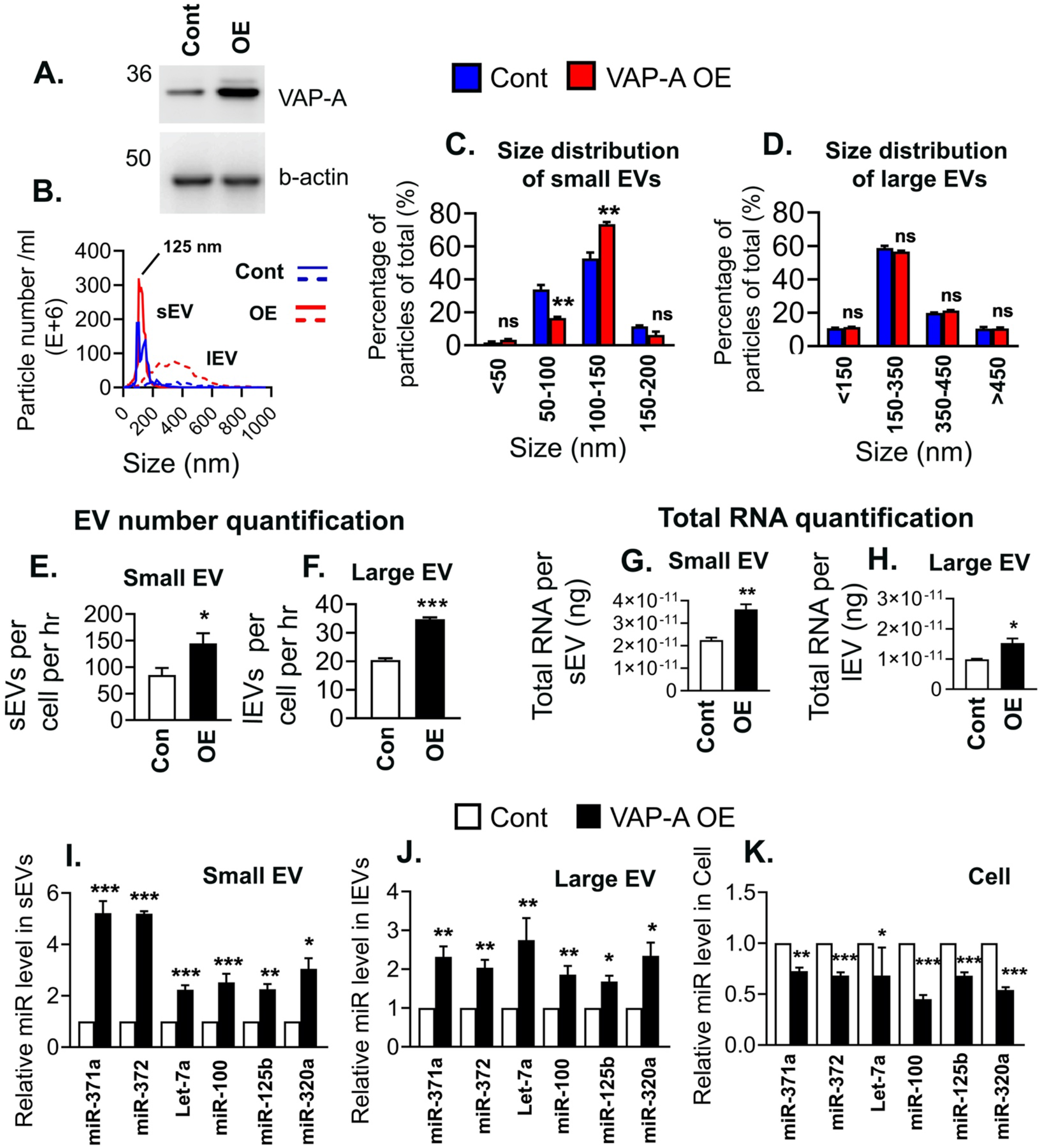
VAP-A overexpression promotes biogenesis of RNA-containing EVs. Related to Figures 1 and 2. (A) Western blot of VAP-A in DKs-8 cells shows overexpression (OE) of VAP-A. Beta actin serves as an endogenous control. (B-D) NTA analysis of size distribution of large (lEV) and small (sEV) EVs purified from control (Cont) and VAP-A OE cells. (B) Representative NTA traces. (C, D) Size distribution of small and large EVs from three independent NTA experiments. (E and F) Calculation of small and large EV secretion rate from control and VAP-OE cells based on NTA analysis of purified EVs and known cell number and media conditioning time. From three independent NTA experiments. (G and H) Total RNA concentration (measured by NanoDrop (A260)) per EV (measured by NTA) for EVs purified from control and VAP-A OE DKs-8 cells. (I-K) Relative levels of specific miRNAs in small and large EVs purified from control (Cont) and VAP-A OE (OE) DKs-8 cells and their parental cells. Data from three independent experiments. Data were plotted as Mean ± S.E.M. *p<0.05, **p<0.01, ***p<0.001.

**Figure S5:**
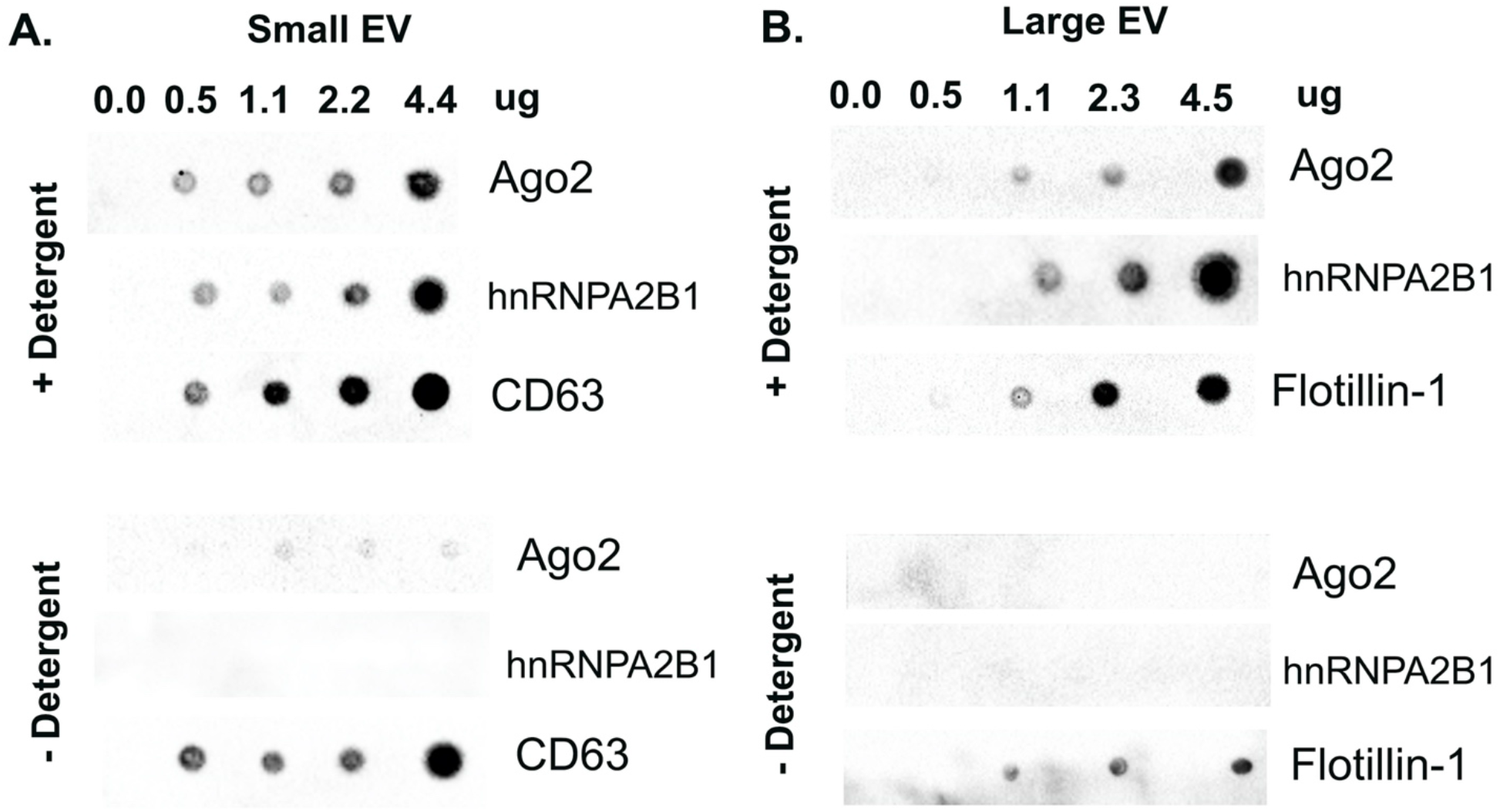
RBPs are present on the inside of small and large EVs. Related to Figure 2. (A and B) Different concentrations of small and large EVs from DKs-8 cells were dotted on nitrocellulose membranes and probed with anti-Ago2, anti-hnRNPA2B1, anti-CD63 or anti-flotillin-1 antibodies in the presence (+Detergent) or absence (-Detergent) of 0.1% Tween-20 as shown. Representative of three independent experiments.

**Figure S6:**
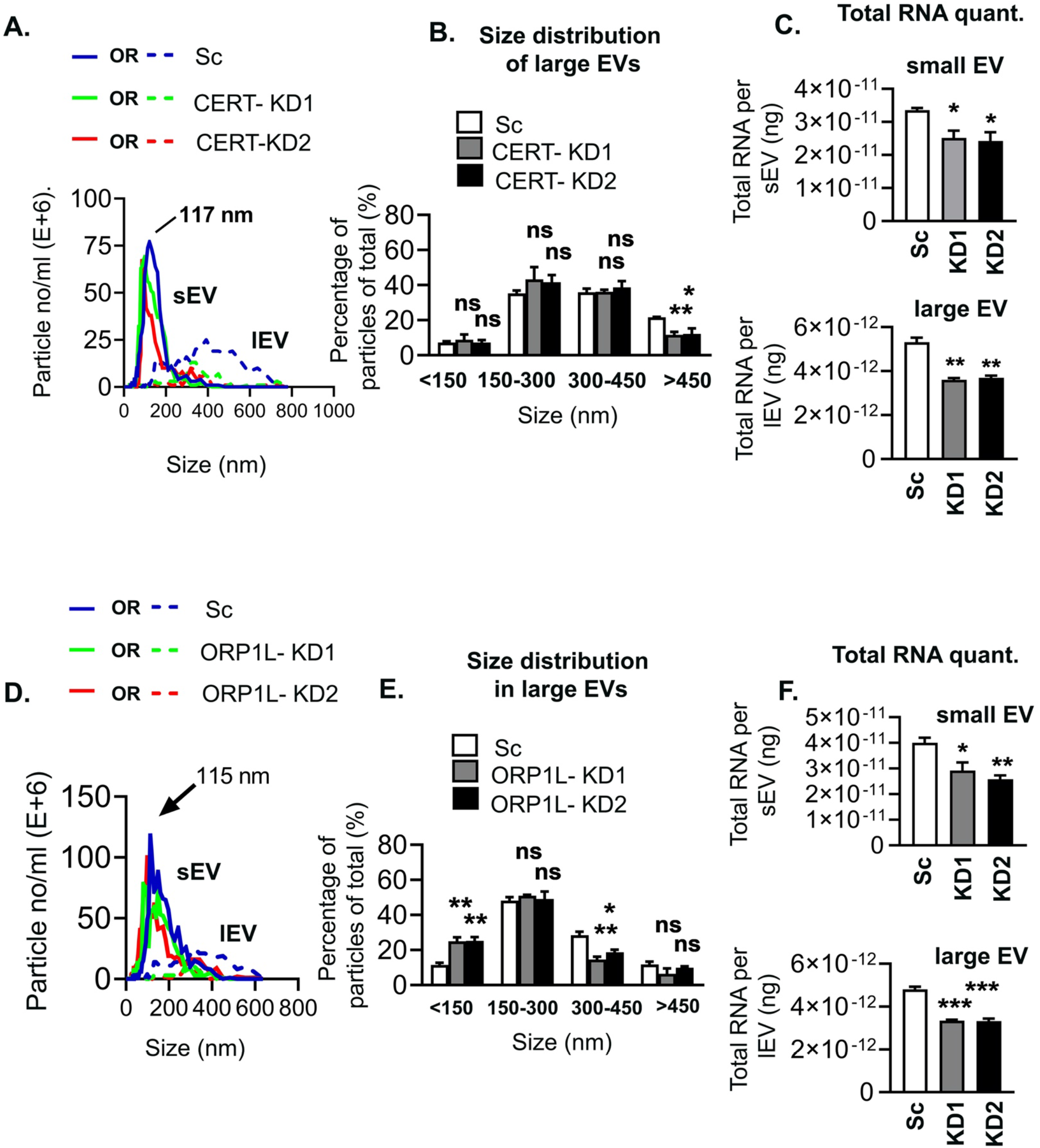
Analysis of small and large EV size and total RNA content as a consequence of CERT or ORP1L KD. Related to Figure 6. (A) Nanoparticle traces of small and large EVs purified from control (Sc) and CERT-KD DKs-8 cells. (B) Size analysis of large EVs purified from control and CERT-KD cells. Data from three independent NTA experiments and plotted as percentage of total large EV number. (C) Total RNA quantity per small and large EV measured by A260 with NanoDrop. Data from three independent experiments. (D) Nanoparticle traces of small and large EVs purified from control (Sc) and ORP1L-KD DKs-8 cells. (E) Size analysis of large EVs purified from control and ORP1L-KD cells. Data from three independent NTA experiments and plotted as percentage of total large EV number. (F) Total RNA quantity per small and large EV measured by A260 with NanoDrop. Data from three independent experiments. Data plotted as Mean ± S.E.M. *p<0.05, **p<0.01, ***p<0.001.

Supplementary Datasheet Legends

Datasheet 1: microRNA data from small RNA-Seq data for control, VAP-A KD1, VAP-A KD2 cells, small EVs and large EVs.

Datasheet 2: snoRNA data from small RNA-Seq data for control, VAP-A KD1, VAP-A KD2 cells, small EVs and large EVs.

Datasheet 3: tRNA data from small RNA-Seq data for control, VAP-A KD1, VAP-A KD2 cells, small EVs and large EVs.

Datasheet 4: Untargeted lipid mass spectrometry data for control, VAP-A KD1, VAP-A KD2 cells, small EVs and large EVs.

